# Human Neonatal MR1T Cells Have Diverse TCR Usage, are Less Cytotoxic and are Unable to Respond to Many Common Childhood Pathogens

**DOI:** 10.1101/2025.03.17.643805

**Authors:** Dylan Kain, Wael Awad, GW McElfresh, Meghan Cansler, Gwendolyn M. Swarbrick, Kean Chan Yew Poa, Conor McNeice, Gregory Boggy, Katherine Rott, Megan D. Null, David M. Lewinsohn, Jamie Rossjohn, Benjamin N. Bimber, Deborah A. Lewinsohn

## Abstract

Neonatal sepsis is a leading cause of childhood mortality. Understanding immune cell development can inform strategies to combat this. MR1-restricted T (MR1T) cells can be defined by their recognition of small molecules derived from microbes, self, and drug and drug-like molecules, presented by the MHC class 1-related molecule (MR1). In healthy adults, the majority of MR1T cells express an invariant α-chain; TRAV1-2/TRAJ33/12/20 and are referred to as mucosal-associated invariant T (MAIT) cells. Neonatal MR1T cells isolated from cord blood (CB) demonstrate more diversity in MR1T TCR usage, with the majority of MR1-5-OP-RU-tetramer(+) cells being TRAV1-2(-). To better understand this diversity, we performed single-cell-RNA-seq/TCR-seq (scRNA-seq/scTCR-seq) on MR1-5-OP-RU-tetramer(+) cells from CB (n=5) and adult participants (n=5). CB-derived MR1T cells demonstrate a less cytotoxic/pro-inflammatory phenotype, and a more diverse TCR repertoire. A panel of CB and adult MAIT and TRAV1-2(-) MR1T cell clones were generated, and CB-derived clones were unable to recognize several common riboflavin-producing childhood pathogens (*S. aureus, S. pneumoniae, M. tuberculosis*). Biochemical and structural investigation of one CB MAIT TCR (CB964 A2; TRAV1-2/TRBV6-2) showed a reduction in binding affinity toward the canonical MR1-antigen, 5-OP-RU, compared to adult MAIT TCRs that correlated with differences in β-chain contribution in the TCR-MR1 interface. Overall, this data shows that CB MAIT and TRAV1-2(-) MR1T cells, express a diverse TCR repertoire, a more restricted childhood pathogen recognition profile and diminished cytotoxic and pro-inflammatory capacity. Understanding this diversity, along with the functional ability of TRAV1-2(-) MR1T cells, could provide insight into increased neonatal susceptibility to infections.

## Introduction

Neonatal sepsis is one of the leading causes of death in children, accounting for an estimated 400,000-700,000 deaths annually.^1^ Tuberculosis (TB), caused by *M. tuberculosis* (Mtb), is another leading cause of death in children, accounting for nearly a quarter million deaths a year.^2^ Neonates are particularly vulnerable to infections, including Mtb, in part due to a naïve and underdeveloped immune system. Understanding the immune system response in early life, and the way in which different immune cells develop can inform vaccine strategies to provide protection in early life from a variety of infections, including TB.

MR1-restricted T (MR1T) cells are defined by their recognition of small molecules including microbial,^3–7^ self,^8,9^ and drug and drug-like molecules^10^ presented by the monomorphic MHC class 1-related molecule (MR1). In adults, the majority of MR1T cells, known as mucosal associated invariant T (MAIT) cells, expresses an invariant TCR α-chain (TRAV1-2/TRAJ33/20/12 in humans), paired with a limited array of Vβ segments,^11^ and express high levels of CD161^12–14^ and CD26.^14^ MR1T cell populations which express TCRα-chains other than TRAV1-2 have been described.^6,15^ Here, we define MAIT cells as a subset of MR1T cells that are TRAV1-2(+) and MR1T cells that express TCRα-chains other than TRAV1-2 as TRAV 1-2 (-) MR1T cells. Riboflavin metabolites are dominant MR1 ligands,^7,16^ and thus MAIT cells recognize riboflavin producing organisms^3,4^ and have been shown in mouse models to be important in immune defense against many common childhood pathogens including *Klebsiella pneumoniae,*^17^ *E. coli,*^4^ and *S. pneumoniae*.^18^ TRAV1-2(-) MR1T cells can respond to a broader array of microbial ligands, including recognition of microorganisms that cannot produce riboflavin.^6^ TRAV1-2(-) MR1T cells also recognize non-microbial ligands and may play additional roles in cancer immunology, auto-immunity and other immune driven diseases.^7,9,19,20^

While TRAV1-2(-) MR1T cells are rare in healthy adult populations, they are the majority of MR1T cells in cord blood (CB) samples.^21^ By 10-weeks of age, however, infant MAIT cells are present in similar frequencies as in adult participants.^21^ Furthermore, infant MAIT cells, were able to produce TNF in response to *Mycobacterium smegmatis* at a similar level to adult MAIT cells, while in CB there were fewer MAIT cells that produced TNF.^21^ The rapid shift from abundant TRAV1-2(-) MR1T cells at birth, to more functional MAIT cells by 10-weeks of age, suggests rapid expansion of MAIT cells early in life, possibly drive by microbial exposures, such as those from the microbiota.^22^

It remains unknown the degree to which the higher frequency of naïve, less functional MR1T cells in CB contribute to the high susceptibility to bacterial pathogens, including Mtb, in the first few months of life. To explore this, as well as the diversity of TCRs in neonatal MR1T cells, we performed combined single-cell-RNA and TCR-sequencing (scGEX-seq/scTCR-seq) on MR1-5-(2-oxopropylideneamino)-D-ribitylaminouracil (MR1-5-OP-RU)-tetramer positive sorted MR1T cells from 5 CB and 5 adult participants. We show that unstimulated MAIT and TRAV1-2(-) MR1T cells from CB have less pro-inflammatory and cytotoxic gene expression profiles compared to adults, as well as significantly more diversity in their TCRs. Both TRAV1-2(+), and TRAV1-2(-), MR1T cell clones isolated from CB retain these features. In addition, none of the CB-derived MR1T clones were able to recognize common childhood pathogens: *S. aureus, S. pneumoniae,* or *M. tuberculosis.* Further, the binding and structural investigations of the CB MAIT TCR clone highlights the importance of the β-chain in MR1-restricted antigen recognition. Collectively, these findings shed light onto the way in which naïve MR1T cells in CB are less capable of host defense to bacterial and mycobacterial pathogens and therefore may contribute to increased risk of these infections in early life.

## Results

### CB MAIT and TRAV1-2(-) MR1T cells are less cytotoxic and pro-inflammatory

To investigate the phenotype and function of MR1T cells in CB, we performed scRNA-seq/scTCR-seq on MR1-5-OP-RU-tetramer(+) cells from CB (n=5) as well as adult controls (n=5). The experiment included 5 oligo-conjugated antibodies for cell surface protein expression at single-cell resolution. After quality control and filtering steps, there were 2,200 adult and 1,513 CB MR1T cells included in the analysis. CB-derived MR1T cells were transcriptionally distinct from adult MR1T cells (Figure 1A). Unsupervised clustering resulted in 9 distinct clusters (Figure 1B) with each cluster predominantly made up of cells from either adult or CB participants (Figure 1C). Cell surface protein expression was different between CB and adult participants, with most adult cells being MAIT cells, expressing not only TRAV1-2 but also CD8, CD26, CD161, while CB cells were predominantly TRAV1-2, CD161, and CD26 negative and more likely to express CD4. (Figure 1D). In CB, CD4+ cells clustered separately from CD8+ cells, whereas in adults, rare CD4+ cells were dispersed throughout the UMAP. Adult MR1T cells also showed increased expression of pro-inflammatory and cytotoxic genes, (*TNF, CCL4, GNLY, GZMA, PRF1*), indicating a more effector-differentiated phenotype (Figure 1E).

**Figure 1:**
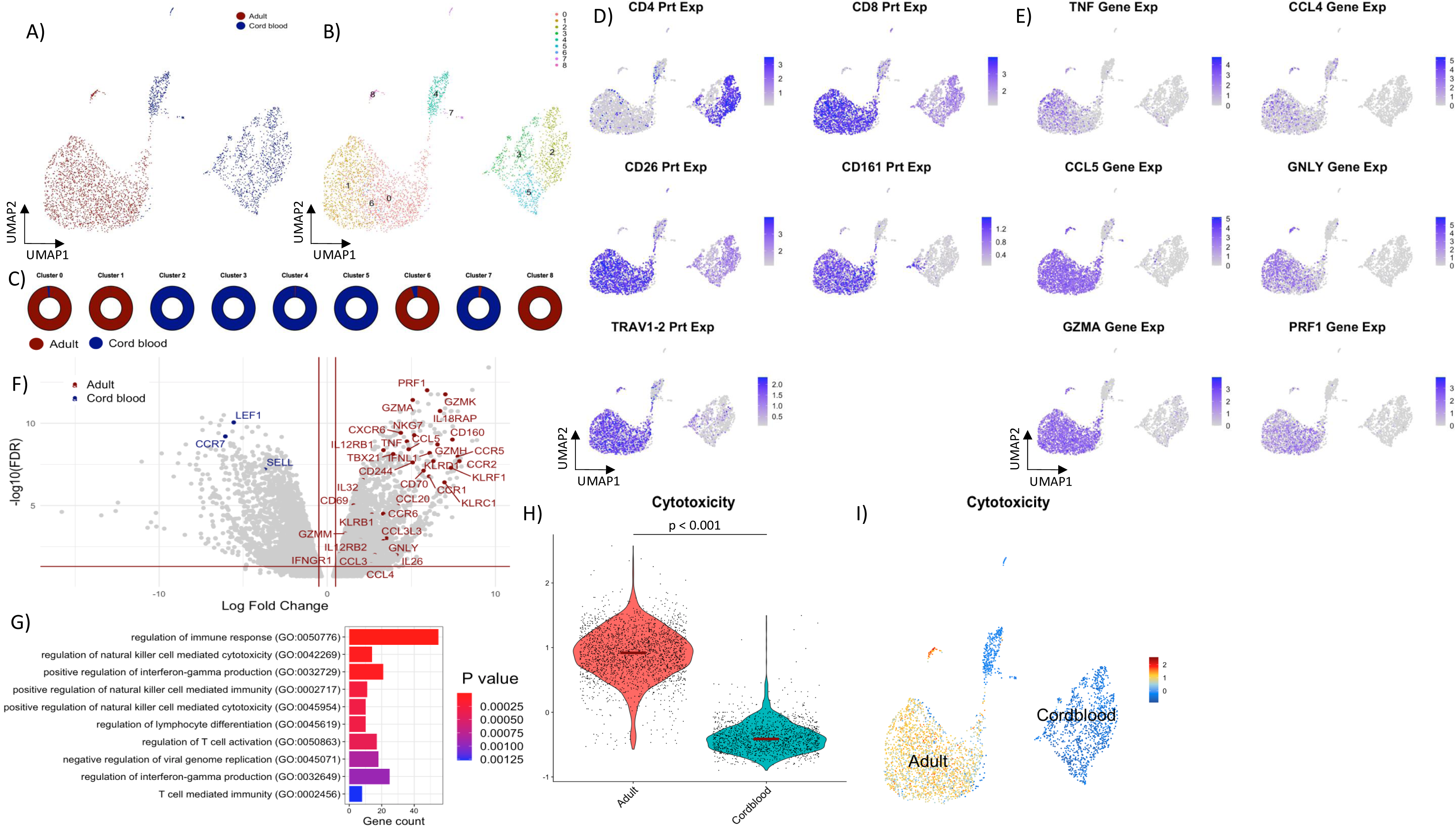
CB MR1T cells have a distinct, less cytotoxic gene expression profile compared to adult MR1T cells. a) Dimensionality reduction of all MR1/5-OP-RU sorted MR1T cells from 5 adult donors and 5 CB donors (n = 2,200 adult MR1T cells and 1,513 CB MR1T cells). b) Unsupervised clustering of all MR1T cells c) Percent of each cluster that is made up by each donor type (adult vs CB) d) Cell surface protein expression by CITE-seq staining. e) Gene expression of common inflammatory/cytotoxic genes f) Volcano plot of differentially expressed genes in CB vs adult MR1T cells. Horizontal red line represents Bonferroni-adjusted p-value of 0.05 and vertical red line represents log fold change of -0.5 or 0.5. g) Top 10 upregulated gene ontology pathways in adult MR1T cells compared to CB MR1T cells. h,i) Cytotoxicity score (*PRF1, GNLY, NKG7, GZMA, GZMB, GZMH, GZMK* and *GZMM*) for adult compared to CB MR1T cells. p < 0.0001 using Wilcoxon test.

Differential gene expression between CB and adult MR1T cells yielded 3083 upregulated genes in adults and 3157 upregulated genes in CB cells (Figure 1F, Supplemental Table 1). Notably, adult cells differentially expressed many pro-inflammatory and cytotoxic genes (*PRF1, GZMA, GZMH, GZMM, GZMK, GNLY, CCL3, CCL4, CCR5, CCR2, CCR1, TNF, NKG7, IL32, IL26*), meanwhile, CB cells had increased expression of naïve markers (*LEF1, CCR7, SELL*). Top upregulated pathways in adult MR1T cells included several pro-inflammatory and activation pathways (Figure 1G). To illustrate these differences, we scored cells for enrichment of a gene module associated with cytotoxic differentiation, comprised of *PRF1, GNLY, NKG7, GZMA, GZMB, GMZK, GZMM,* using the UCell package.^23^ Unstimulated adult cells were more cytotoxic than CB MR1T cells (Figure 1H,I). In CB participants the majority of TRAV1-2(-) cells were CD4-positive and CD8-negative, whereas in adult participants the majority of cells were CD8-positive irrespective of TRAV1-2 status (Supplemental Figure 1).

### CB MR1T cells have far more diversity of their TCR repertoire

Next, we examined the diversity of TCR usage in adult and CB MR1T cells, finding significantly more diversity in CB samples (Figure 2). Surface TRAV1-2 staining was much lower for CB cells (Figure 1D) and this was confirmed through scTCR-seq, demonstrating that TRAV1-2 represents less than a quarter of all TCRs used (Figure 2A,D). Furthermore, while canonical TRAJ chains, TRAJ33/20/12, are also the most common TRAJ chains used by CB cells, they represent a far smaller subset of total TRAJ compared to adult cells (Figure 2B,E). CB TRBV chains also showed more diversity compared to adult MR1T cells (Figure 2C,F). This diversity is quantified using the Shannon diversity index, showing significantly more diversity in CB TCRs (Figure 2G-I). Furthermore, adult participants were found to have more expanded clonotypes, with 19.1% (107/560) of cells having at least one other cell with an identical CDR3, whereas only 0.1% (1/959) of CB cells were expanded clonotypes (Supplemental Figure 2).

**Figure 2:**
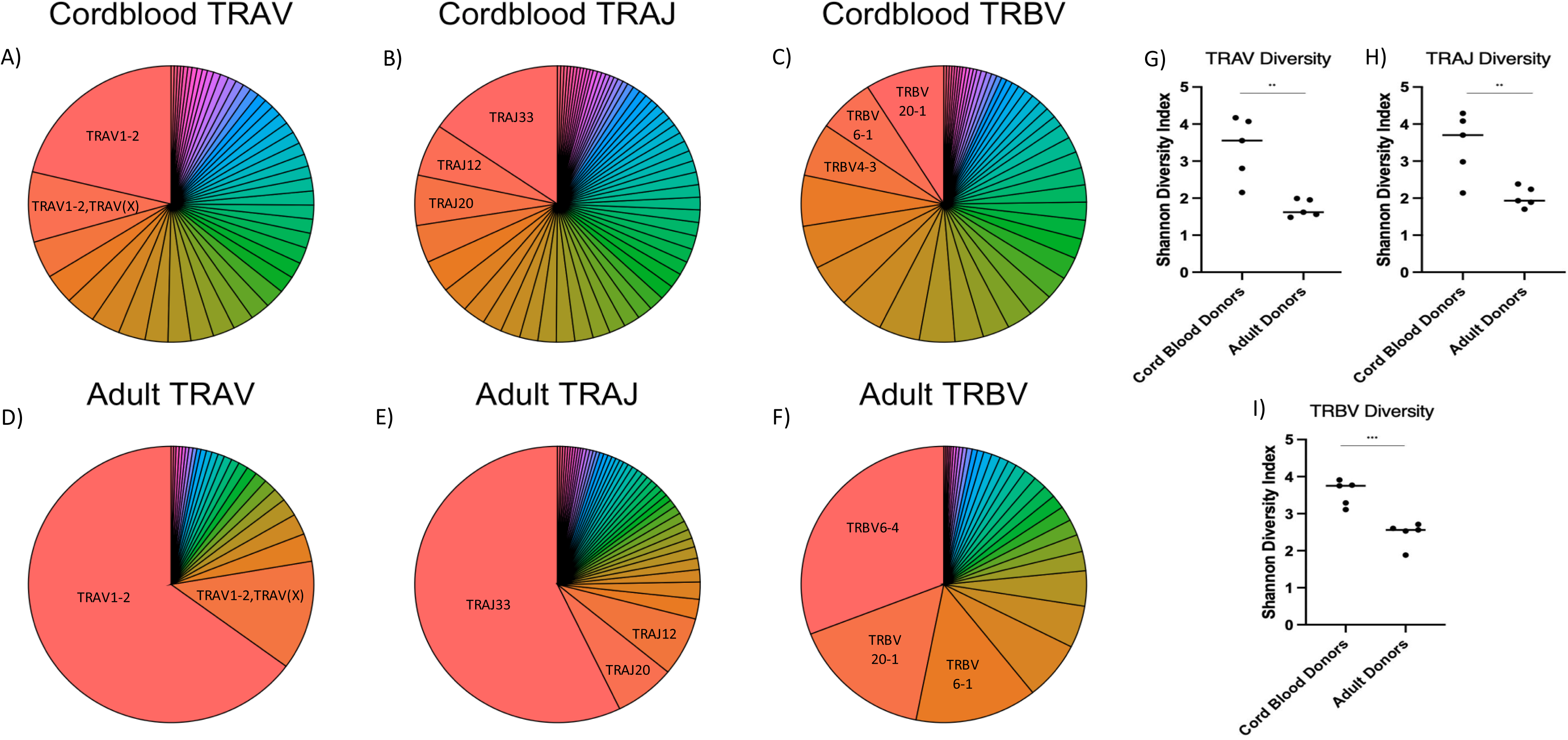
CB MR1T cells have a much more diverse TCR repertoire compared to adult MR1T cells. a-f) CB and adult MR1T cell TRAV, TRAJ and TRBV repertoire by TCR sequencing. TRAV1-2,TRAV(X) represents cells with dual expression of alpha chains, one being TRAV1-2 and TRAV(X) representing any TRAV1-2(-). g-i) Shannon diversity index of TRAV, TRAJ and TRBV repertoire of CB and adult MR1T cells. Unpaired t-test with p-value < 0.01 denoted by ** and p-value < 0.001 denoted by ***

Previous work has shown that dual TCR-alpha expression can confound MAIT cell TCR interpretation.^24^ To confirm that this would not be the driving force in TCR interpretation, we examined cell surface protein expression of TRAV1-2 utilizing CITE-seq staining in CB participants, comparing cells with TRAV1-2 gene expression compared to those without (Figure 3). TRAV1-2 gene expressing cells had much higher cell surface expression of TRAV1-2, suggesting dual TCR-alpha expression would not significantly affect our subsequent analysis. Given the high frequency of TRAV1-2(-) MR1T cells in the CB participants, we examined if TRAV usage would impact TRAJ or TRBV pairings. Utilizing TCR sequences from CB participants, we compared TRAJ usage for TRAV1-2(+) MAIT cells compared to TRAV1-2(-) cells (Figure 4A). TRAV1-2(+) cells had significantly higher pairing with TRAJ33, whereas TRAV1-2(-) cells had significantly higher pairing preference with non-canonical TRAJs (non-TRAJ33/20/12). This suggests TRAV1-2 usage influences TRAJ usage in MR1T cells. We next explored if there were pairing preference differences in MAIT cells from CB compared to adult participants. Examining only TRAV1-2(+) cells, we compared TRAJ usage between adult and CB participants (Figure 4B). Even among MAIT cells, adult participants had significantly higher canonical TRAJ33 pairings compared to CB participants. Adult participants’ MAIT cells also had significantly higher pairing preference for TRBV20-1 (Figure 4C).

**Figure 3:**
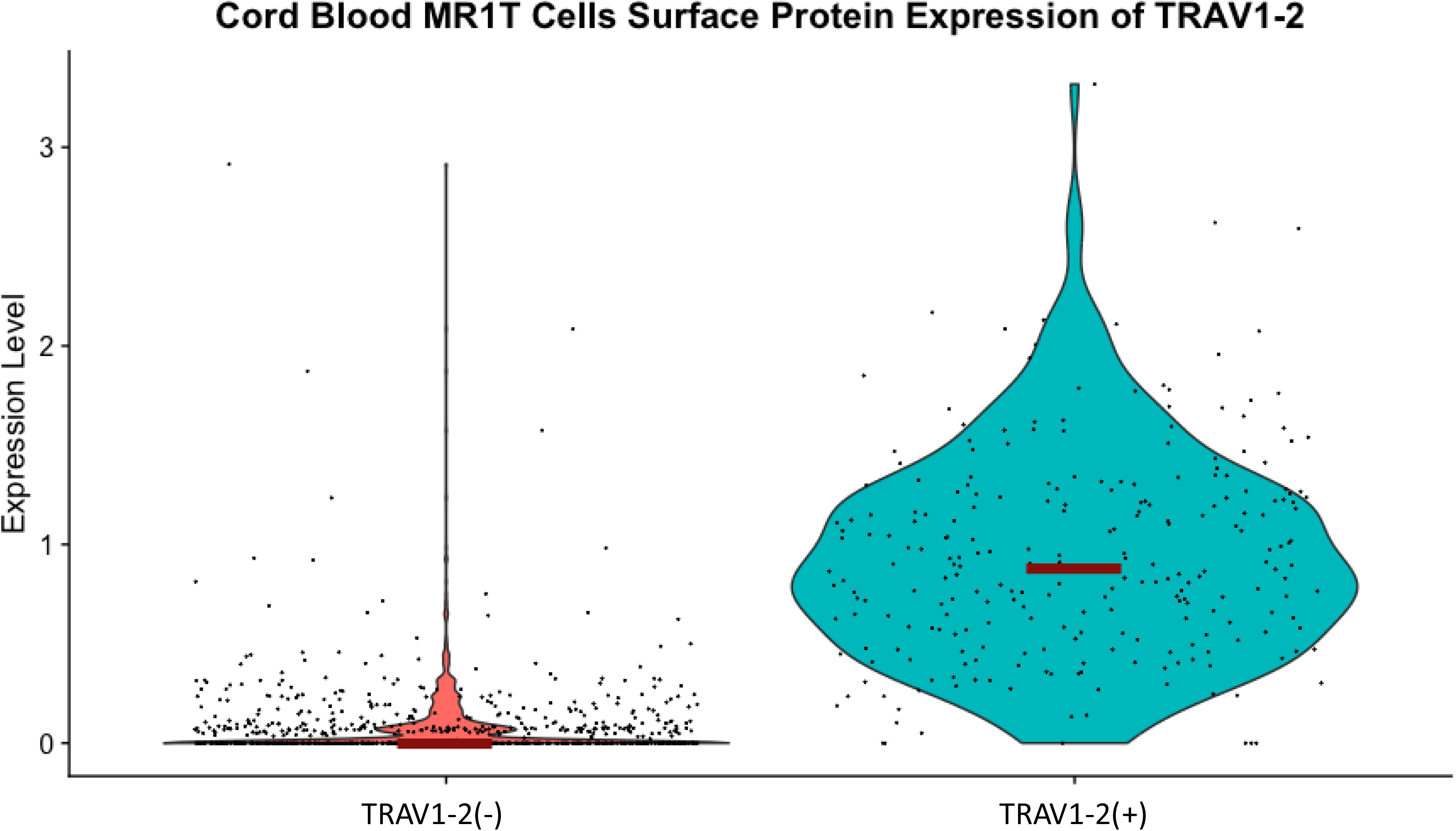
TRAV1-2 cell surface protein expression matches TRAV1-2 gene expression for CB MR1T cells. CB MR1T cells were sorted into TRAV1-2(+) or TRAV1-2(-) based on gene expression, and violin plot of cell surface protein expression is displayed based on CITE-seq staining (cell surface protein expression) for TRAV1-2 (TCR Vα7.2)

**Figure 4:**
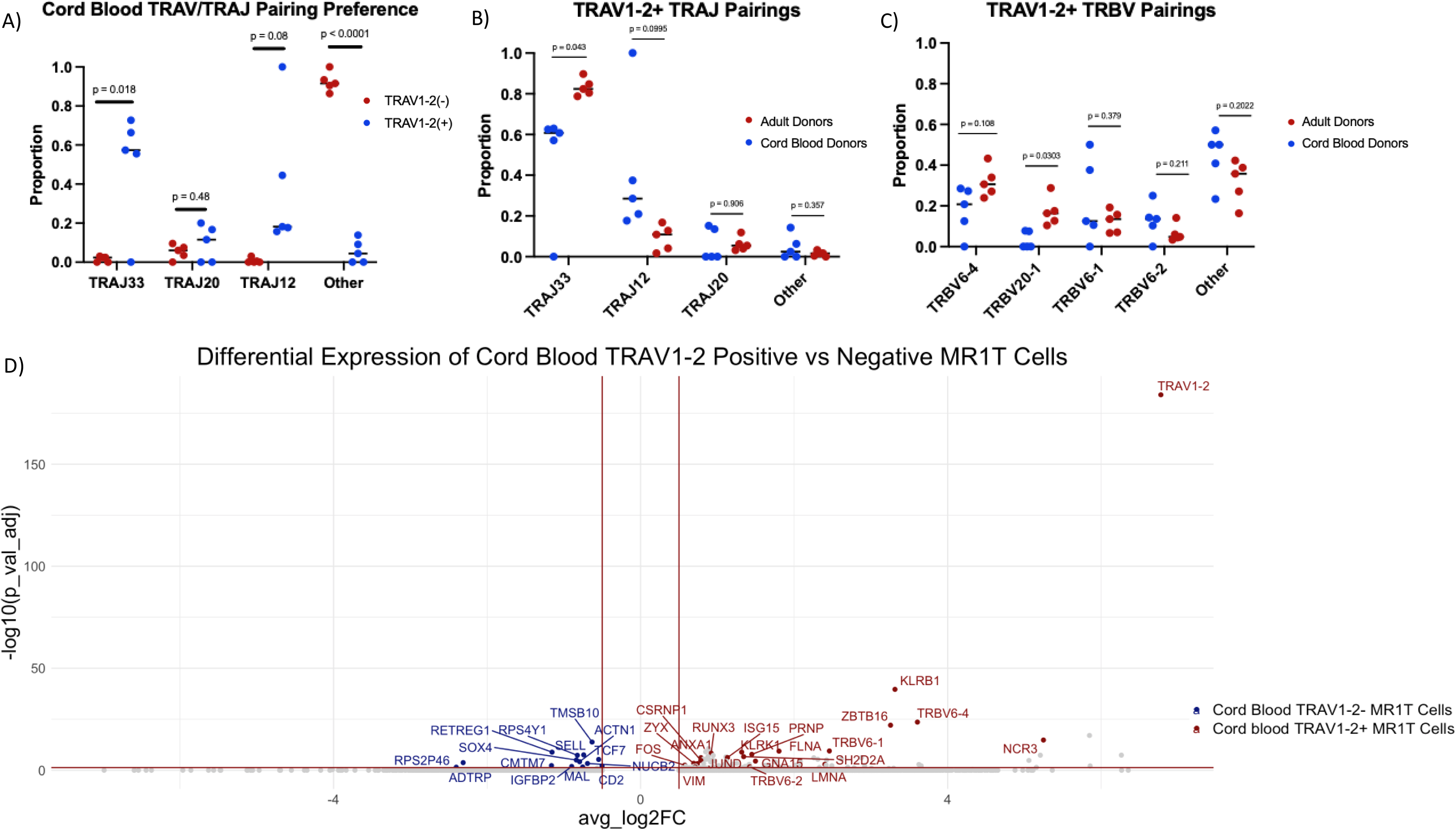
Pairing preferences more diverse for CB MR1T cells. a) Pairing preferences of CB MR1T cells based on gene expression of TRAV1-2(+) vs TRAV1-2(-). Unpaired t-test. b) TRAJ pairing preference for TRAV1-2(+) MR1T cells from adult and CB donors. Unpaired t-test. c) TRBV pairing preference for TRAV1-2(+) MR1T cells from adult and CB donors. Unpaired t-test. d) Volcano plot for differential gene expression of CB MR1T cells expressing TRAV1-2(+) or TRAV1-2(-). Labeled genes were greater than 0.5 or less than -0.5 log fold change and had a Bonferroni-adjusted p-value of < 0.05.

### Unstimulated CB MAIT cells are more mature than CB TRAV1-2(-) MR1T cells

Given the increased gene expression of pro-inflammatory and cytotoxic genes in adult-derived MR1T cells, and the much higher percentage of MAIT cells in adults, we hypothesized that MAIT cells in CB would also have higher expression of pro-inflammatory and cytotoxic gene expression. Therefore, we examined subsets of CB-derived MAIT and TRAV1-2(-) MR1T cells and performed differential gene expression between TRAV1-2(+) and TRAV1-2(-) cell populations. This yielded 15 genes that were significantly upregulated in TRAV1-2(-) cells and 80 genes that were significantly upregulated in TRAV1-2-(+) cells (Figure 4D and Supplemental Table 2). As expected, *TRAV1-2* was the most increased gene among TRAV1-2(+) cells. TRBV genes, *TRBV6-1/TRBV6-4,* were also significantly upregulated in TRAV1-2(+) cells, they were more likely to express TRBV6-1, TRBV6-2 or TRBV6-4. Other genes that were significantly upregulated in TRAV1-2(+) cells include effector function genes (*KLRB1, KLRK1, NCR3, ZBTB16*),^13^ genes associated with T-cell-activation (*LMNA, ANXA1, PRNP, CSNP1*),^25–27^ genes involved in cell proliferation and cell signaling (*SH2D2D, JUND, FOS, GNA15*),^28–30^ interferon associated genes (*ISG15, ZYX*)^31^ and genes involved in tissue residency or tissue trafficking (*RUNX3, VIM, FLNA*).^25,32,33^ Although there are limited significantly upregulated genes in TRAV1-2(-) cells, notably they do upregulate *SELL*, (encoding for CD62L/L-selectin), whose expression is generally high on naïve T cells.^34^

### Unstimulated adult MR1T cells are more cytotoxic than CB MR1T cells

Since there is an expansion of MAIT cells over the first year of life, one possible explanation for the increased cytotoxicity expression profile demonstrated by adult as compared to CB MR1T cells (Figure 1G), is the increased proportions of MAIT cells in adults compared to neonates. To explore this, we compared the cytotoxicity scores of the TRAV1-2(+) cell subset from CB to adult participants (Figure 5A,B). Adult MAIT cells from adult participants had much higher cytotoxicity scores compared to those from CB participants, suggesting that even the CB MAIT cell subset of MR1T cells demonstrate decreased cytotoxicity when unstimulated compared to adult MAIT cells. This is also reflected in the differentially expressed genes between CB MAIT and adult MAIT cells (Figure 5C, Supplemental Table 3), wherein adult MAIT cells upregulated cytotoxicity and effector genes (*GZMA, GZMB, GZMK, GZMM, GNLY, PRF1, NKG7, CST7, KLRD1, KLRG1, CCL4, CCL5, IL32*) and downregulate naïve and repair genes (*LEF1, SELL, CCR7, TCF7, VEGFA, VEGFB*). This suggests that CB MAIT cells continue to mature in after birth in the periphery.

**Figure 5:**
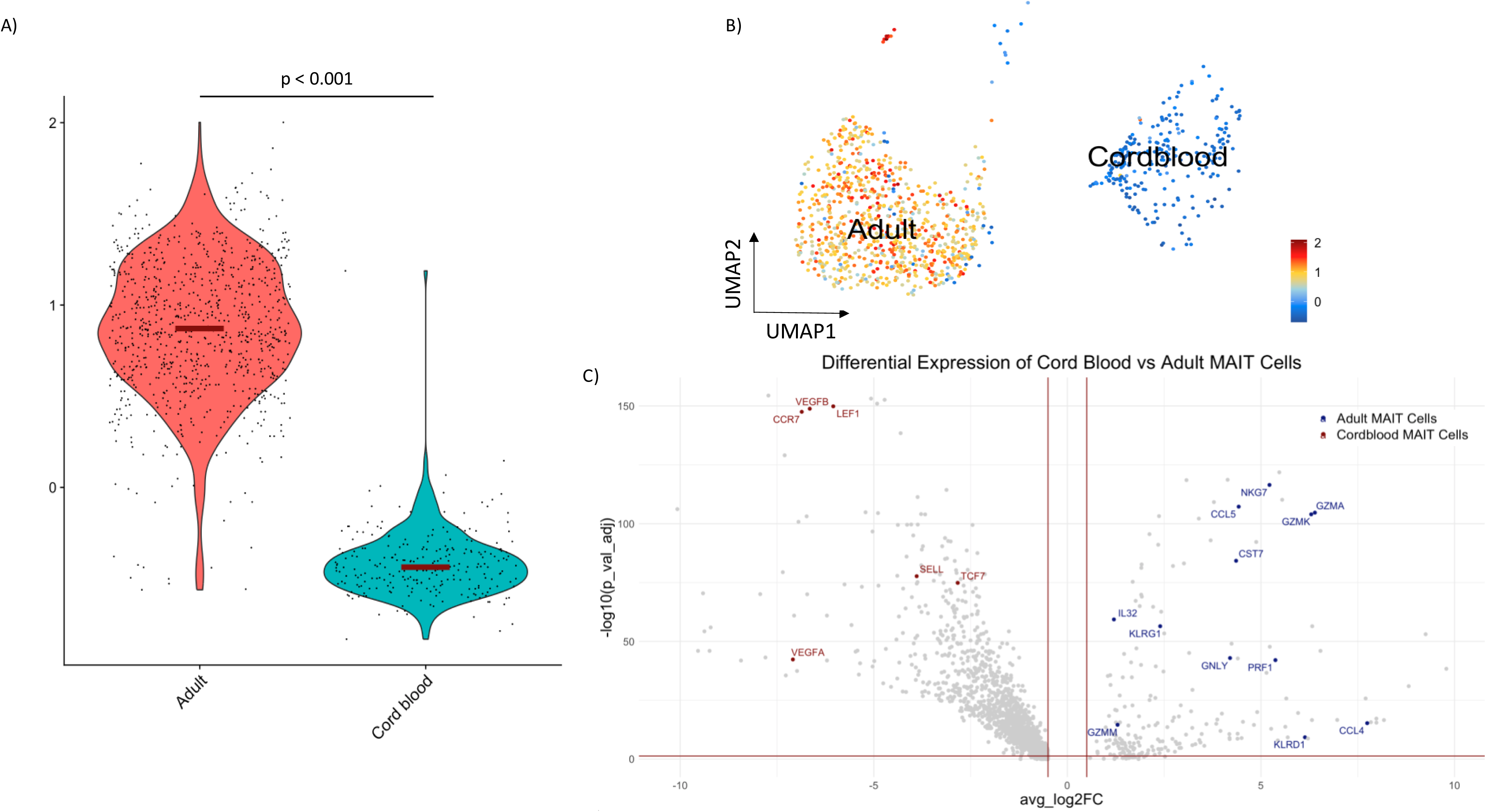
TRAV1-2(+) adult MAIT cells are more cytotoxic than TRAV1-2(+) CB MAIT cells. a-b) Cells were sorted for TRAV1-2(+) MR1T cells and then cytotoxicity score (*PRF1, GNLY, NKG7, GZMA, GZMB, GZMH, GZMK* and *GZMM*) was assessed for adult compared to CB MR1T cells. p-value < 0.0001 calculated using Wilcoxon test. c) Volcano plot for differential gene expression of CB MAIT vs Adult MAIT. Labeled genes were greater than 0.5 or less than -0.5 log fold change and had a Bonferroni-adjusted p-value of < 0.05.

### Increased phenotypic diversity of CB MR1T cell clones as compared to adult MR1T cell clones

MR1T cell clones were isolated by limited dilution assay (LDA) of MR1-5-OP-RU-tetramer(+) cells derived from one adult participant (D719) and one CBMC specimen (CB964). Herein, we characterize two CB TRAV1-2(-) MR1T clones (CB964 A1/A4), one CB MAIT clone (CB964 A2) and three adult MAIT clones (D719 A2/C1/C2). Given the rarity of MAIT and TRAV1-2(-) MR1T cells at birth and the high rate of TRAV1-2(-) MR1T cells, we first validated that clones that were generated represented the diversity of TCR usage seen in CB participants. Figure 6A shows the TCR diversity of the *ex-vivo* MR1-5-OP-RU-tetramer(+) cell population from CB964 and Figure 6B shows the TCR usage of the three clones from this participant, illustrating that T cell clones isolated using LDA are representative of the diverse TCR repertoire of this participant. Figure 6C and 6D show the tetramer staining and TCR usage of each of the MAIT and TRAV1-2(-) CB clones and each of the three adult MAIT clones. As expected, all MR1T cell clones retained strong MR1/5-OP-RU staining. MR1/6-formylpterin (MR1/6-FP) did not stain the MAIT cell clones, while both TRAV1-2(-) CB clones did demonstrate MR1/6-FP staining. CB964 A1 uniformly expressed CD8 and variably expressed CD4 (Supplemental Figure 3A). CB964 A2 and A4 expressed both CD8 and CD4 (Supplemental Figure 3B,C). Each of the adult MAIT clones utilized TRAV1-2/TRAJ33 and was CD8(+)CD4(-) (Supplemental Figure 3D-F).

**Figure 6:**
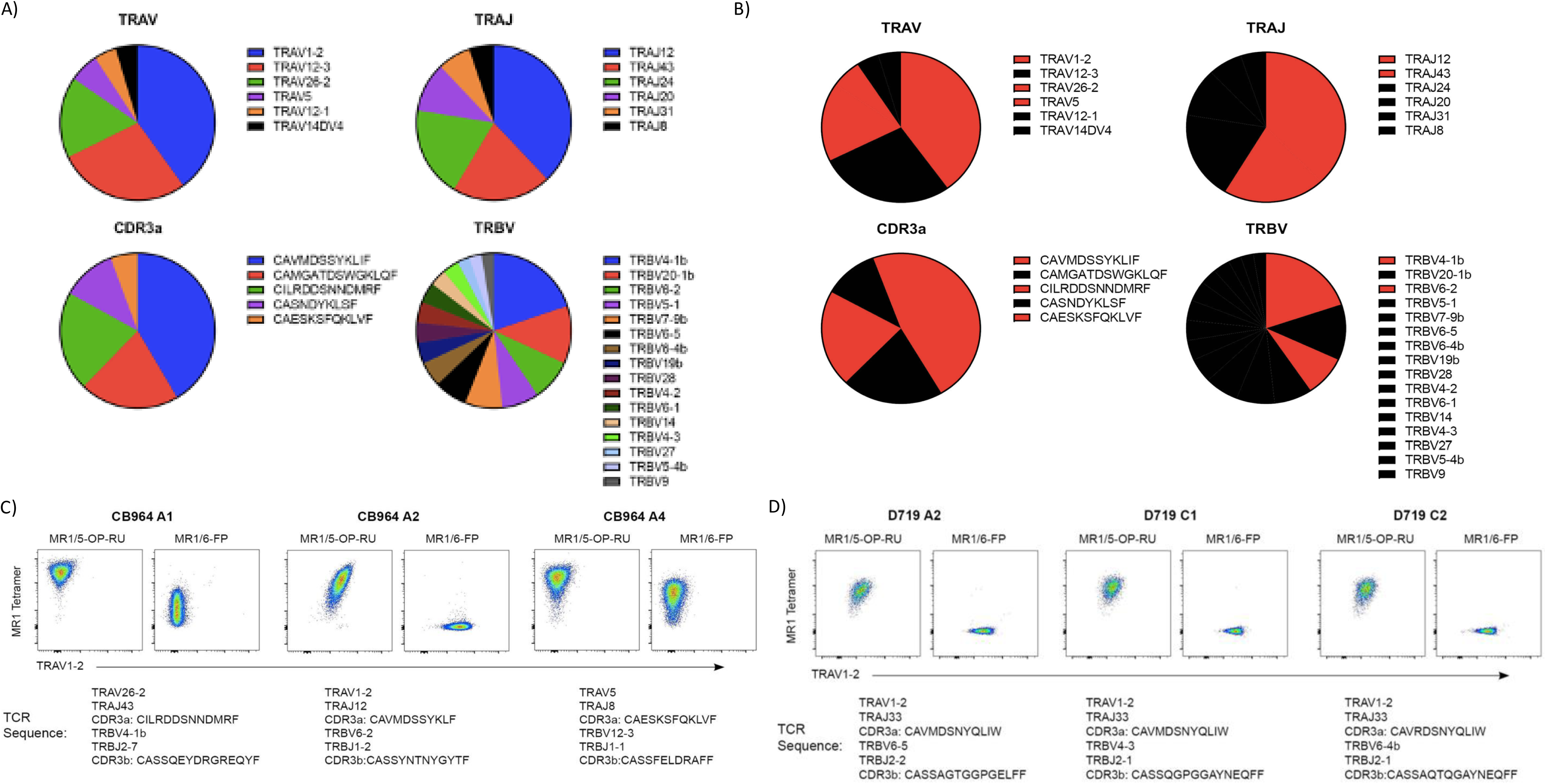
MR1T cell generated clones can represent the diversity of TCR repertoire of CB donors. a) TCR repertoire of donor CB964 from which CB MR1T cell clones were generated. b) Red represents TCR repertoire from donor CB964 for which MR1T cell clones were generated. c) MR1/5-OP-RU and 6-FP tetramer staining by flow cytometry of CB MR1T cell clones along with TCR sequence obtained by single-cell sequencing below for each of CB964 A1, CB964 A2 and CB964 A4. d) MR1/5-OP-RU and 6-FP tetramer staining by flow cytometry of adult MR1T cell clones along with TCR sequence obtained by single-cell sequencing below for each of D719 A2, D719 C1 and D719 C2.

### CB MR1T cell clones demonstrate different unstimulated and stimulated gene expression profiles from adult MR1T cell clones

To test the functional response of CB MR1T cells compared to adults, PMA/ionomycin stimulated and unstimulated clones were analyzed via single-cell-sequencing. Similar to adult MAIT cell clones, CB MAIT and TRAV1-2(-) MR1T cell clones were able to produce typical pro-inflammatory and cytotoxic cytokines following stimulation (Figure 7A, Supplemental Figure 4). Differential gene expression comparing CB and adult clones shows many significantly differentially expressed genes both unstimulated and following stimulation (Figure 7B, Supplemental Table 4,5). Notable trends include unstimulated CB MR1T cell clones have increased expression of naïve genes (*FUOM, SELL, CCR7, IL7R, LEF1, RGS10*), along with increased expression of many chemokines (*CCL3, CCL4, CCL23, CCL18, CCL1, CCl22, CCL24, CCL8, CCL13, CCL7, CCL2, CXCL9, CXCL12, CXCL3, CXCL16*) and increased type I interferon genes (*IFI30, IFI27, IFI44L, IFIT3, IFIT1, IFI6, IFI27L1, ISG15, MX2, IRF4, IRF9, IRF8, HERC3*) as compared to unstimulated adult MAIT cells clones. Unstimulated adult MAIT cell clones show increased expression of cytotoxic genes (*GZMH, PRF1, LTB, FASLG*) along with glycolysis genes (*PGAM5, ALDOC, PFKP, PGK1, ENO1, ENO2, GAPDH, PKM*), associated with effector phenotype in MAIT cells as compared to unstimulated CB MR1T cell clones.^35^ Following stimulation, CB MR1T cells continued to have increased chemokines and type I interferon genes expression relative to adult MAIT cell clones, while adult MAIT clones demonstrate activation markers (*CD8A, TNFRSF14, NR4A3, FASLG, LTB, RGCC, CST7*), pro-inflammatory cytokines (*IL26, IL32, TNF*), glycolysis genes (*PFKP, ENO2, PGK1, PKM, HK2, ALDOC*) and exhausted T-cell genes (*LAG3, CD160, CTLA4, CD244*) as compared to CB MR1T cell clones.

**Figure 7:**
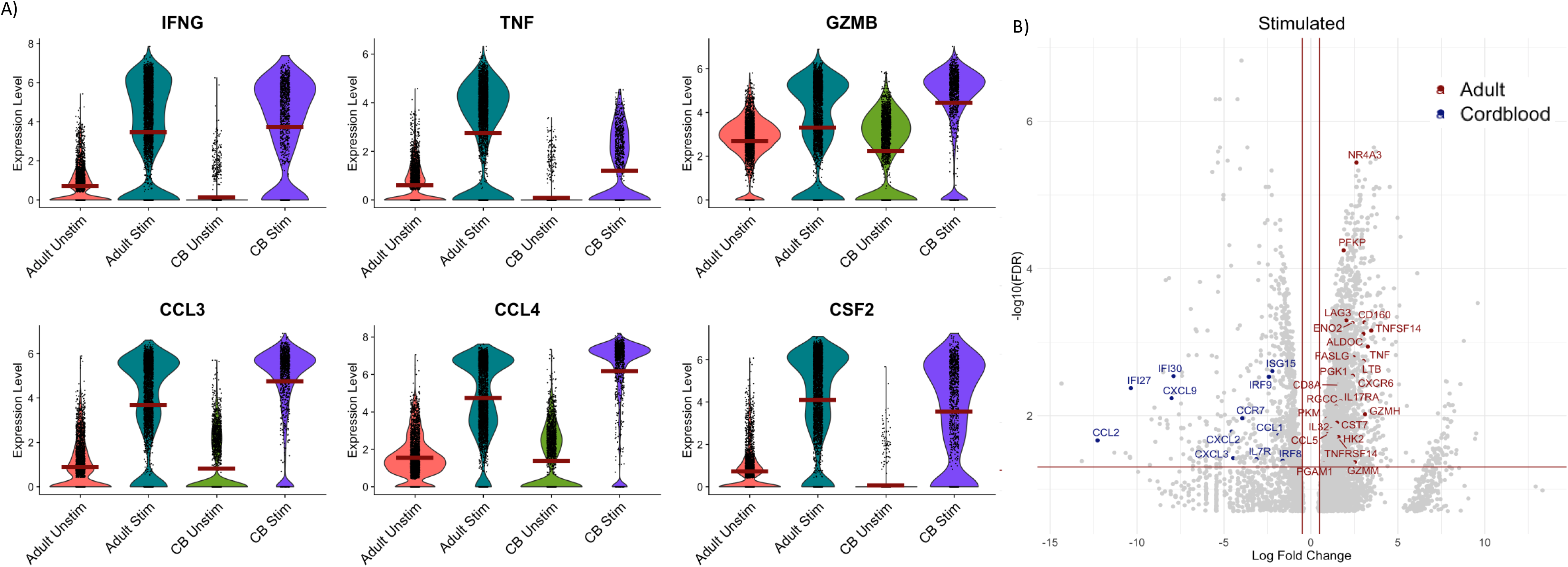
CB MR1T cell clones can produce typical inflammatory/cytotoxic cytokines by gene expression, but have distinct gene expression profiles from adult MR1T cell clones. a) Violin plot of gene expression of common inflammatory/cytotoxic genes from stimulated (PMA/Ionomycin) and unstimulated MR1T cell clones from adult compared to CB donors b) Volcano plot of differentially expressed genes of adult compared to CB MR1T cell clones when stimulated with PMA/Ionomycin. Horizontal red line represents Bonferroni-adjusted p-value of 0.05, and vertical red lines represent log fold change of 0.5 or -0.5.

### CB MR1T cell clones demonstrate differential recognition of childhood pathogens

We next tested CB MR1T cell clones against a panel of common childhood pathogens (*E. coli, S. aureus, S. pneumoniae)* as well as various mycobacteria (*M. tuberculosis* auxotroph; mc^2^6206 (H37Rv ΔpanCD ΔleuCD),^36^ *M. smegmatis*),and mycobacterial MR1T mycobacterial ligands (photolumazine 1 [PL1], deazalumazine [DZ])^5^ and compared this to an adult MAIT clone well studied in our lab; D426 G11.^6^ All of these bacteria/mycobacteria are riboflavin producers and thus should activate MR1T cells.

Each clone had a very different pattern of microbe and antigen recognition by IFNγ ELISpot (Figure 8A-H) and this recognition was blocked by anti-MR1-antibody, demonstrating MR1-dependence. While adult-derived G11, demonstrated strong responses to *E. coli, S. aureus,* and *S. pneumoniae*, none of the CB MR1T clones recognized *S. aureus* or *S. pneumoniae* and recognized *E. coli* less well (CB964 A1/A2), or not at all (CB964 A4). Regarding mycobacteria, the adult-derived clone, G11, responded to Mtb auxotroph, while none of the CB MR1T cells recognized Mtb auxotroph. G11 demonstrated a robust response to *M. smegmatis*, such that MR1 blocking was not apparent until using lower concentrations of the microbe (Supplemental Figure 6) and selectively recognized DZ but not PL1 as previously described.^5^ CB964 A1 similarly recognized *M. smegmatis* and DZ but not PL1. By comparison CB964 A2 robustly responded to *M. smegmatis* in an MR1-dependent but did not recognize PL1 or DZ, while CB964 A4 did not recognize *M. smegmatis* despite strongly recognizing PL1 and DZ. Thus, these three CB MR1T cell clones each expressing different TRAV’s, demonstrate differential recognition of *M. smegmatis* and known mycobacterial MR1T ligands. Finally, all clones were tested against human monocyte-derived dendritic cells (DC) in the absence of microbes or antigens (background media control), and none demonstrated self-reactivity (data not shown).

**Figure 8:**
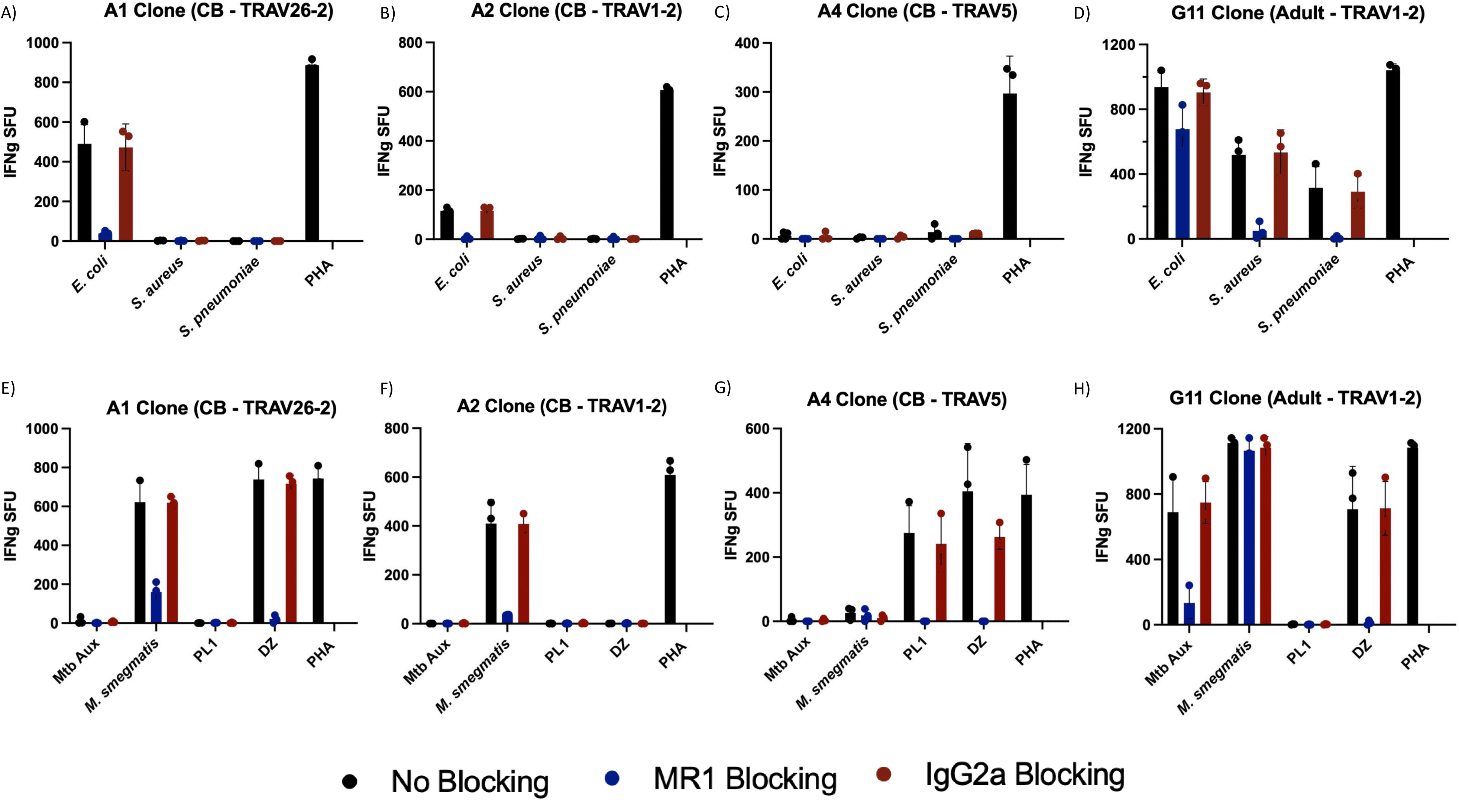
CB and adult MR1T cell clones have distinct and diverse microbial recognition pattern. a-d) IFNγ ELISPOT of CB MR1T cell clones (A1, A2 and A4) and adult MR1T cell clone (G11) response to common childhood bacterial pathogens. Responses without blocking, anti-MR1 blocking antibody or isotype control blocking antibody are represented. Each clone was tested against DCs without microbes or antigens added, and this background was subtracted from IFNγ SFU shown. e-h) IFNγ ELISPOT of CB MR1T cell clones (A1, A2 and A4) and adult MR1T cell clone (G11) response to Mycobacteria and Mycobacterial antigens. Responses without blocking, anti-MR1 blocking antibody or isotype control blocking antibody are represented. SFU = spot forming units. PHA = phytohaemagglutin P. PL1 = photolumazine I. DZ = deazalumazine.

### A CB MAIT TCR displayed a different affinity for MR1 compared to two adult MAIT TCRs

To establish how adult and CB MAIT TCRs bind to MR1, three soluble extracellular domains of the adult and CB MAIT TCRs were expressed as inclusion bodies in *Escherichia coli*, refolded into their native conformation and purified.^37^ This included two adult MAIT TCRs: A-F7 (TRAV1-2-TRBV6-1), #6 (TRAV1-2-TRBV6-4),^38^ and CB964 A2 TCR clone (TRAV1-2/TRBV6-2) (Figure 9A). Next, Surface Plasmon Resonance (SPR) experiments were performed to examine the specificities and determine the steady state binding affinities (K_D_) of A-F7, #6, and CB964 A2 TCRs against MR1-5-OP-RU or MR1-Ac-6-FP complexes (Figure 9B). As previously reported, the adult MAIT A-F7 and #6 TCRs bind only to MR1-5-OP-RU with high affinity (K_D_ ∼2-4 µM),^37^ while the CB964 A2 TCR exhibited a roughly 5-fold lower affinity of K_D_ ∼18 µM to MR1-5-OP-RU and did not bind to MR1-Ac-6-FP (Figure 9B). As such, the CB MAIT TCR tested was riboflavin-based ligand reactive yet exhibited lower affinity towards MR1-5-OP-RU.

**Figure 9:**
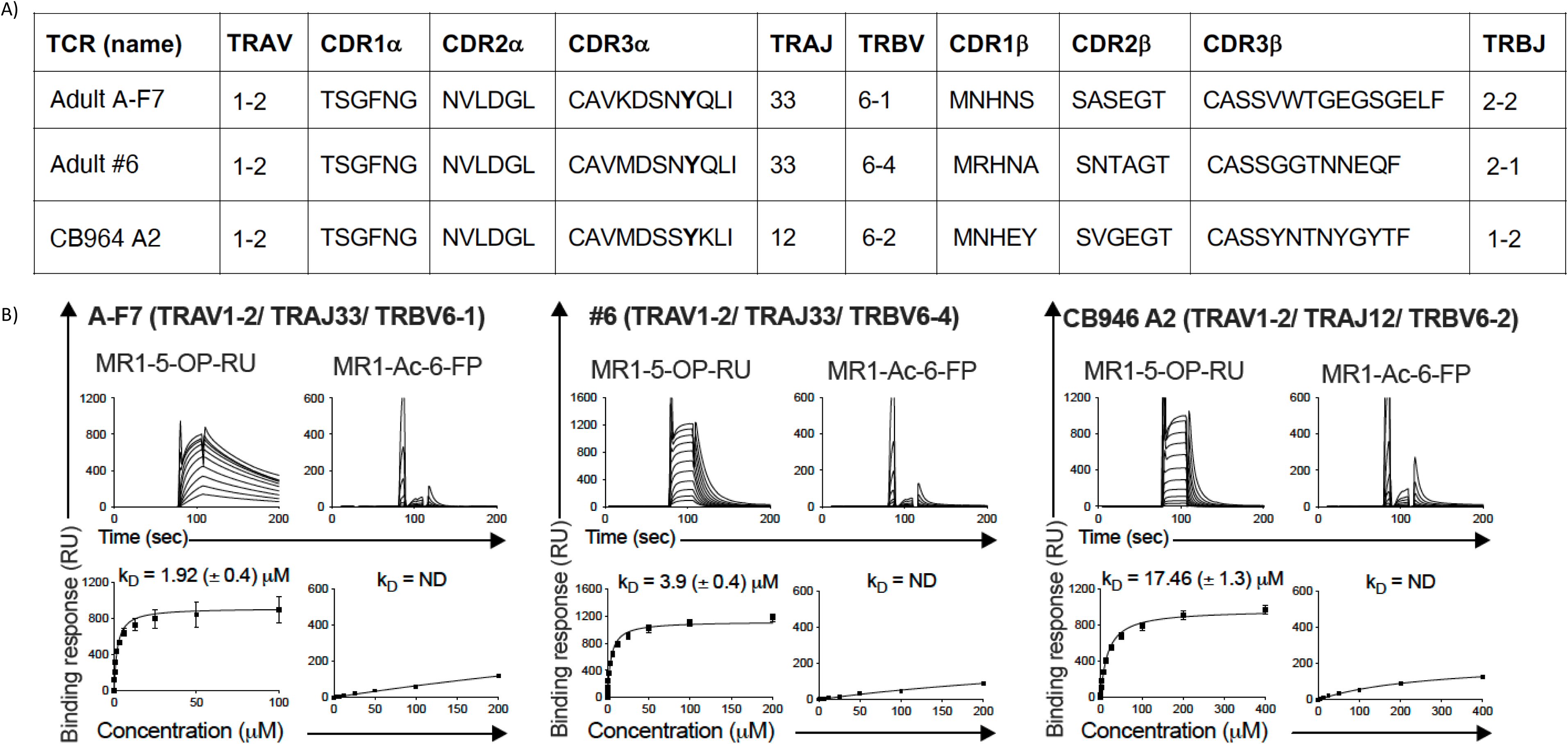
Steady-state affinity measurements of soluble TCRs for MR1-antigen complexes. a) TCR αβ gene names and CDR amino acid sequences of the TCR cell clones. b) The affinity of TCR-MR1-Ag interactions were determined using SPR, by measuring the binding of various concentrations of soluble adult MAIT TCRs of A-F7 (TRAV1-2/TRBV6-1; *left panel*) and #6 (TRAV1-2/TRBV6-4; *middle panel*), as well as cord CB946 A2 TCR clone (TRAV1-2/TRBV6-2; *right panel*) against human MR1 refolded with 5-OP-RU and Ac-6-FP antigens. The SPR runs were conducted as duplicate in two independent experiments using different batches of proteins. Experiments were conducted with serial dilutions of the TCRs. The SPR sensograms (*upper panels*), equilibrium curves (*lower panels*) and steady state *K*_D_ values (µM) were prepared in GraphPad Prism 10. Error bars represent the mean and SD from technical replicates (two independent experiments). ND, not determined. RU, response unit

### Overview of CB MAIT TCR-MR1-5-OP-RU ternary complex

We next determined the crystal structure of the CB964 A2 (TRAV1-2/TRBV6-2) TCR-MR1-5-OP-RU ternary complex at 2.5Å resolution and compared it with the published adult MAIT A-F7 (TRAV1-2-TRBV6-1) TCR-MR1-5-OP-RU structure^37,39^ (Figure 10A-D, Supplemental Table 6). The 5-OP-RU ligand in the CB964 A2 TCR-MR1-5-OP-RU complex was sequestered within the A′ pocket and formed a Schiff-base covalent bond with MR1-Lys43, consistent with all published adult MAIT TCR-MR1-5-OP-RU complexes.^40^ The CB964 A2 TCR docked orthogonally atop the MR1 antigen binding cleft, with the TCR α- and β-chains positioned atop the α2- and α1-helices of MR1, respectively (Figure 10A,B). The buried surface area (BSA) at the interface between CB964 A2 TCR and MR1-5-OP-RU was approximately ∼1030Å^2^, a value that falls close to the range of BSA of adult MAIT TCR-MR1-5-OP-RU complexes (1060-1200Å^2^). Here, the centre of gravity of the CB964 A2 α and β-chains were similar to the corresponding chains of the A-F7 TCR (Figure 10E-F). While the α- and β-chains of adult MAIT TCRs contributed almost equally to the BSA of the interfaces of most TCR–MR1-Ag complexes, the α- and β-chains of the CB964 A2 TCR contributed to ∼54.4 % and 45.6 % respectively (Figure 8B,D), with this difference being attributed to slightly differing MR1 docking modality of the CB964 A2 TCR β-chain (Figure 10E-F). Specifically, the CB964 A2 TCR was slightly rotated in comparison to adult MAIT TCRs, which positioned the β-chain slightly away from MR1 and reduced the contribution of CB964 A2 TCR β-chain to the interface with MR1. Consequently, the CB964 A2 TCR adopted a moderately different footprints on MR1 compared to adult TCRs (Figure 10B,D).

**Figure 10:**
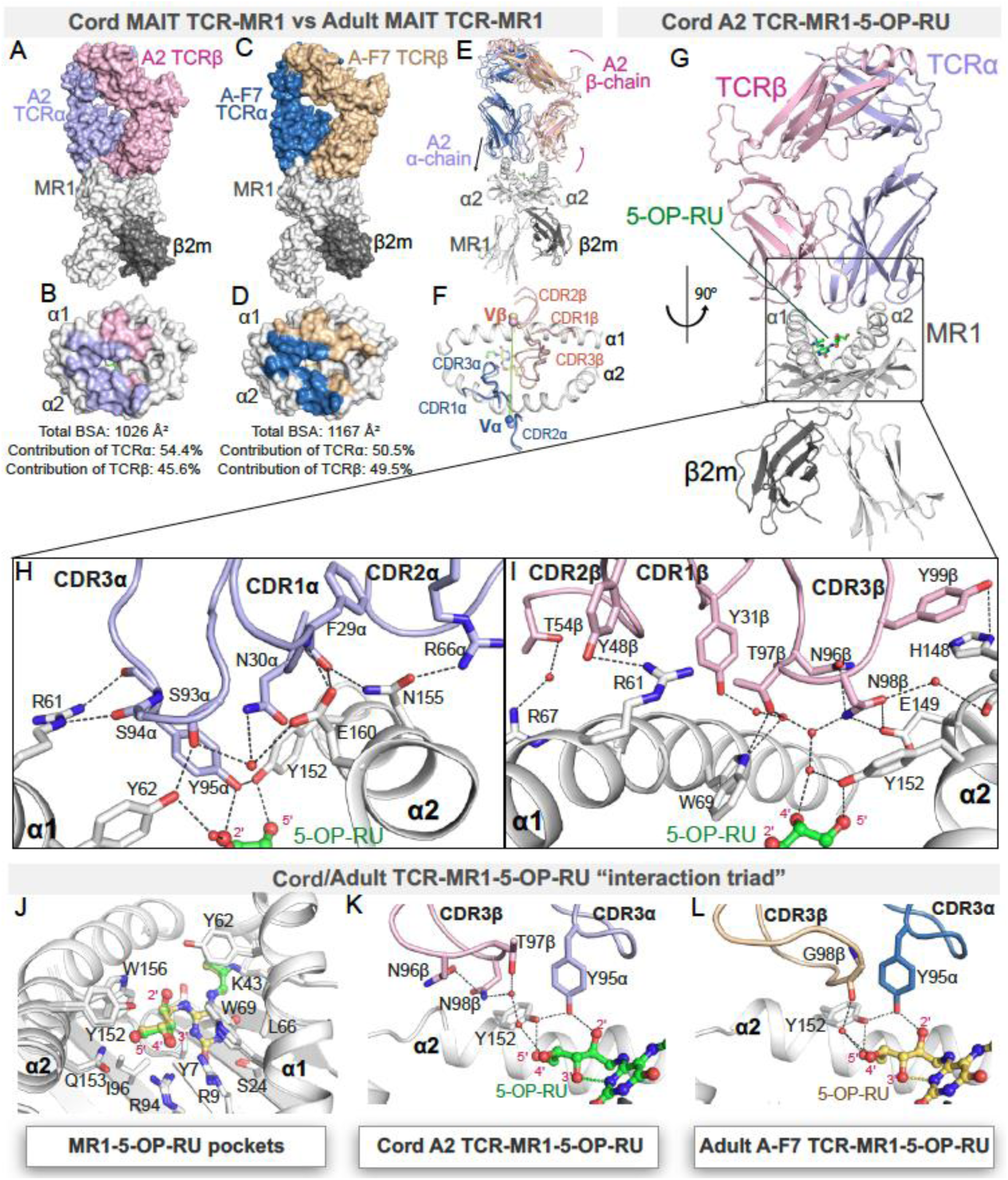
Structural comparison of ternary complexes of cord A2 and adult AF-7 typical MAIT TCRs with MR1-5-OP-RU. a-d) Crystal structures of ternary complexes of (a-b) cord A2 MAIT (TRAV1-2/TRBV6-2) TCR-MR1-5-OP-RU and (c-d) adult MAIT A-F7 (TRAV1-2/TRBV6-1) TCR-MR1-5-OP-RU (PDB ID: 6PUC). The top panels (*a and c)* are molecular surface representations of the ternary complexes; the lower panels (b and d) illustrate the respective TCR footprints on the molecular surface of MR1-5-OP-RU. The MR1 and β2-microglobulin molecules are colored white and dark grey, respectively and 5-OP-RU is presented as green and yellow sticks in A2 and A-F7 ternary structures, respectively. A2 TCRα, lavender; A2 TCRβ, light-pink; A-F7 TCRα, sky-blue; A-F7 TCRβ, wheat. The atomic footprint is colored according to the TCR chain making contact. e) Comparison of cord A2 TCR ternary complex relative to the adult A-F7 TCR docking positions. Arrows illustrate TCR rotation around the center of mass of the MR1, as well as displacement of the β-chain along the MR1 binding cleft. f) Superposition of the CDR loops of cord MAIT A2 and adult MAIT A-F7 TCRs sitting atop MR1. The center of mass of the respective TRAV and TRBV variable domains are shown as spheres in the same colors as the respective variable domains. g) Cartoon representation of the ternary structure of A2 TCR-MR1-5-OP-RU. h-i) Interactions between (h) the CDRα loops, (i) CDRβ loops of the cord A2 and MR1 residues. The interacting residues are represented as sticks. Hydrogen bond interactions are represented by black dashes. See also Supplemental Table 6. j) Comparison of the MR1 antigen-binding pocket and the position of MR1 residues in both the cord A2 and adult A-F7 ternary structures. k-l) Interactions of the CDR3α and CDR3β of the cord A2 (k) and adult (l) MAIT TCRs, depicting the ‘interaction triad’ between Tyr152 of MR1, Tyr95α of CDR3α and 5-OP-RU.

### Conserved recognition of riboflavin-derived 5-OP-RU by a CB and adult MAIT TCRs

The TRAV1-2 TCR α-chain docks similarly in both CB964 A2 and adult A-F7 TCRs, contributing to ∼560Å^2^ and 590Å^2^ to the BSA of the interface of CB964 A2 TCR-MR1 and adult TCR-MR1 complexes, respectively. The TRAV1-2-TRAJ12 α-chain of the CB964 A2 TCR comprises a conserved set of TRAV1-2-encoded CDR1α residues; Gly28α, Phe29α and Asn30α and CDR2α residues; Val50α and Leu51α. These residues mediate conserved polar and van der Waals (vdw) interactions with residues of MR1 α2-helix, such as His148, Asn155 and Glu160 (Figure 10G,H, Supplemental Table 7). Additionally, the TRAJ12-encoded CDR3α motif residues, ^93^Ser-Ser-Tyr-Lys^96^, of the CB964 A2 TCR forms conserved polar interactions with MR1-His58, Arg61, Tyr62, Tyr152, and Trp156 (Figure 10H). Akin to the adult A-F7 TCR-MR1-5-OP-RU complex, the conserved Tyr95α of the CB964 A2 TCR formed a hydrogen bond with 2′OH group of 5-OP-RU and Tyr152 of MR1 α1-helix, thus forming the *‘interaction triad’* (Figure 10J-L). The positioning of the ribityl tail of 5-OP-RU was conserved in both structures and governed by polar interactions mediated by MR1 residues, Arg9, Arg94, Tyr152 and Gln153 (Figure 10J). Collectively, the CB964 A2 TCR α-chain TRAV1-2 interactions with MR1 were conserved when compared to the published adult MAIT TCR footprints on MR1.^15,38^

The TRBV6-2 β-chain of the CB964 A2 TCR contributed to ∼470Å^2^ of the BSA of TCR-MR1 complex, but the TRBV6-1 β-chain of the adult TCR-MR1 complex was more closely associated with MR1 (BSA∼580Å^2^). Indeed, the TRBV6-1 of the adult A-F7 TCR is oriented closer to the MR1 α1-helix than the CB964 A2 TCR. As a result, it produces more CDRβ-MR1 mediated contacts, which affects their BSA to MR1 (Figure 10B,D,F). Here the CB964 A2 CDR3β loop was essentially the main β-chain participant to the complex interface, followed by CDR2β, whereas the CDR1β loop was not involved in polar interactions with MR1 (Figure 10I). Specifically, the Asn98β and Tyr99β of the CB964 A2 CDR3β loop formed hydrogen bonds with Glu149 and His148 residues of MR1 α2-helix respectively, whereas the Tyr48β of the CDR2β motif formed hydrogen bond with Arg61 and vdw contacts with Arg61 and Gln64 of the of the MR1 α1-helix (Figure 10I, Supplemental Table 7). In accordance with these structural results and SPR affinity data, the CB964 A2 MAIT TCR can recognize riboflavin-derivative-based 5-OP-RU presented by MR1, despite having a weaker interaction with MR1 than adult MAIT TCRs.

## Discussion

Understanding MR1T cell development and early life capacity could help elucidate mechanisms of enhanced susceptibility to infection in neonates and inform strategies to mitigate this risk. Here, we compare *ex-vivo* MR1/5-OP-RU-tetramer(+) cells from CB and adult participants, demonstrating that *ex-vivo* CB-derived MR1T cells exhibit a lower cytotoxicity and inflammatory gene expression profile, accompanied by a much more diverse TCR repertoire. Notably, CB-derived MAIT and TRAV1-2(-) MR1T cell clones failed to respond to several common childhood pathogens (*S. aureus, S. pneumoniae, M. tuberculosis*), despite their riboflavin production. Crystal structure analysis of a CB MAIT TCR-MR1-5-OP-RU complex compared to an adult-derived MAIT TCR revealed that the β-chain binding reduced TCR affinity for an MR1 ribityl ligand. To our knowledge, this represents the first comprehensive transcriptional profile of CB-derived MR1/5-OP-RU-tetramer(+) MR1T cells, including TRAV1-2(-) subsets, and the first detailed analysis of the capacity of these cells to recognize microbes and microbial MR1 ligands. These findings enhance our understanding of MR1T cell development in early life and their role in neonatal immune susceptibility.

It has previously been shown that MAIT cells, defined as TRAV1-2+, CD161 high cells, acquire anti-microbial activity, including the ability to produce IFNγ in utero, prior to the development of the microbiota and thus prior to microbial antigen exposure.^41^ We have previously shown that at birth, the majority of MR1T cells are in fact not MAIT cells and are instead TRAV1-2(-) MR1T cells and that functional MR1T cells as defined by TNF production to *M. smegmatis*, were exclusively contained in the MAIT cell subset.^21^ In addition, MAIT cells rapidly expand in early life, whereas TRAV1-2(-) MR1T cells persist, but remain as memory T cells in adult populations. Taken together, these data support the idea that MR1T cells, as more comprehensively defined by MR1/5-OP-RU-tetramer(+), in CB have diminished capacity to recognize microbes because MAIT cells are relatively infrequent. Our work similarly finds that in CB MAIT cells upregulate genes that suggest they could be more functional compared to TRAV1-2(-) MR1T cells. We extend this work, however, finding that even among CB MAITs there is higher expression of naïve genes, and lower expression of cytotoxicity genes compared to adult MAITs. This includes upregulation of *GNLY*, which has recently been suggested to mark MAIT cells that may be primed for further cytotoxicity.^42^ These findings indicate that at birth MAIT cells are not only less frequent, but also less mature, demonstrating less potential for functional capacity than adult MAIT cells. This maturation over life would therefore most likely happen in the periphery in response to environmental exposures after birth.

Furthermore, our results confirm and extend the prior characterization of human MR1T cell development. CB-derived MR1T cells had significantly higher expression of genes associated with naïve cells and stage 2 MR1T cells than did adult MR1T cells. This is in keeping with previously published work that explored MR1/5-OP-RU-tetramer(+) TRAV 1-2(+) cells, from CB, showing that the majority are stage 2 or early stage 3.^43^ *LEF1* in particular has been shown to be a marker of early stage 3 MAIT cells,^44^ and is also found to be significantly upregulated in CB-derived TRAV1-2(-) MR1T cells and also CB MAIT cells compared to adult MR1T cells. This further supports that immature TRAV1-2(-) MR1T cells and MAIT cells are present in the CB of humans and that extrathymic maturation of these cells occurs after birth.

It has previously been demonstrated that at birth, there is significantly more α-chain diversity in the TCR repertoire of MR1T cells.^21^ We again see this in our data, but also extend this finding, observing significantly more diversity among the entire TCR usage of MR1T cells at birth. Among CB MAIT cells these is increased α-chain diversity as evidenced by decreased canonical TRAV1-2/TRAJ33, TRAV1-2/TRBV6-4 and TRAV1-2/TRBV20-1 pairings. Among TRAV1-2(-) MR1T cells, we observe no skewing of either α- or β-chain usage. This degree of TCR diversity among the TRAV1-2(-) MR1T cells is similar to the that observed among conventional MHC1a-restricted T cells.^45^ We speculate that TRAV1-2(-) MR1T cells may have the capacity to respond to cognate antigenic exposure, albeit the full repertoire of ligands recognized remains to be determined. Therefore, while we find tremendous TCR diversity among MR1T cells at birth, the functional consequences of this remains largely unexplored. Among MAIT cells, the TCR β-chain can contribute to MR1 ligand discrimination.^38,46–48^ Crystal TCR structure from the CB MAIT clone, showed that although it bound 5-OP-RU, it did so with a much lower affinity than would be expected for most MAIT cells, with the α-chain bound 5-OP-RU in a conventional manner, but with less β-chain interaction than is typical. This adds to the previous literature suggesting a role of the β-chain in fine-tuning the MAIT cell antigen response and provides a structural basis for why the β-chain interaction with MR1 is important for TCR recognition of microbial ligands.^38,46–48^

To study the pathogen recognition pattern and functional capacity of CB MR1T cells, we isolated MR1T cell clones from CB. We found that these CB-derived MR1T cell clones are phenotypically and functionally representative of *ex-vivo* CB MR1T cells, demonstrating similar diversity of CD4/CD8 and TCR expression. Furthermore, we observed similar trends of differential gene expression between unstimulated CB clones compared to adult clones, as were seen comparing *ex-vivo* MR1T cells from CB and adult donors. Upon stimulation, we observed that these CB MR1T clones demonstrate pro-inflammatory capacity in a similar manner to adult MR1T clones. In addition, CB MR1T clones upregulated chemokines and ISGs compared to adult clones, and although the small number of clones analyzed limits the ability to generalize these results, it suggests that they may retain their capacity to respond in a TCR-independent manner.

Cloning MR1T cells from CB also allows detailed examination of TCR recognition of MR1 ligands. Despite all CB-derived clones staining for MR1/5-OP-RU-tetramer none of these clones responded to some common riboflavin-producing childhood pathogens; *S. aureus, S. pneumoniae* and *M. tuberculosis*. The reasons for this discrepancy are not entirely clear, but there are several possible explanations. One possibility is that the concentration ligand processed and presented during the infection could be insufficient from these pathogens to activate these clones. Supporting this, we see that the CB-derived CB964 A4 clone, responds to DZ and PL1, both of which are produced by *M. smegmatis,*^5^ but not to live *M. smegmatis*. Another possibility, however, is that the optimal ligands for the TCRs of the CB-derived MR1T cell clones are not produced by these organisms. It’s recently been shown, that for adults, 5-amino-6-D-ribityaminouracil (5-A-RU) is a key precursor for the production of MAIT cell ligands,^49^ but this may not be the case for the diverse TCRs found CB MR1T cells. It is therefore possible that these CB TCRs could recognize 5-OP-RU, but not a sufficiently activating ligand produced by these particular riboflavin-producing organisms. Overall, the lack of response of the CB MR1T cell clones to several common childhood pathogens, suggests that in early life there is a defect in the repertoire of the MR1T cells response to these organisms, that is augmented over time, presumably due to environmental exposures after birth.

It has previously been suggested that TRAV1-2(-) MR1T cells recognize self-antigens, in the absence of microbes.^19^ Here, we find that neither of the two TRAV1-2(-) MR1T clones tested here showed evidence of response to self. Furthermore, we find that one of the TRAV1-2(-) MR1T clones was able to recognize *E. coli*. Therefore, as shown previously,^6^ we find that TRAV1-2(-) MR1T cells can also recognize microbes. As autoreactive TRAV1-2(-) MR1T cells must necessarily survive thymic selection, we postulate that the expansion of these cells must occur later in life, as might also occur with microbially reactive TRAV1-2(-) MR1T cells.

MAIT cells play an important role in defense against many microbial infections, including many common bacterial causes of childhood infections.^4,17^ Our results are consistent with the decreased frequency of functional MR1T cells at birth could play a role in susceptibility to bacterial infection in early life. However, we do also find both MAIT and TRAV1-2(-) MR1T cells at birth that are microbial responsive and have effector abilities when stimulated and could be especially critical in early life prior to the development of classical adaptive immune responses.^50^ The shift in TCR repertoire from CB to adults, suggests that exposures after birth shape these cells to become more functional, and therefore suggests that these cells could be harnessed in early life for the development of novel childhood vaccines.

## Methods

### Study participants

This study was conducted according to the principles expressed in the Declaration of Helsinki. Study participants, protocols, and consent forms were approved by the Institutional Review Board at Oregon Health & Science University (OHSU; IRB00000186). All ethical regulations relevant to human research participants were followed. Umbilical CB was obtained from the placenta of uncomplicated term pregnancies at OHSU in Portland, Oregon, after delivery into citrate CPT tubes (BD). Blood was processed per manufacturer’s instructions within 24 hours of collection and the resulting CB mononuclear cells (CBMC) were cryopreserved. Adult peripheral blood mononuclear cells (PBMC) were obtained via leukapheresis from healthy adult participants and cryopreserved. Adult participants lacked any detectable responses to Mtb immunodominant antigens, CFP-10 and ESAT-6 and were therefore assumed to be Mtb uninfected.

### Single-cell sequencing of MR1T cells and MR1T cell clones

The following reagents were obtained through the NIH Tetramer Core Facility: MR1-5-OP-RU tetramer and MR1/6-FP tetramer.^39^ Identification of CB and adult MR1T cells with MR1 tetramers was described previously.^11,21^ CBMC or PBMC were thawed in the presence of DNase, resuspended in 10% heat inactivated human serum with RPMI (Lonza) at a concentration of 2x10^7^/ml. CBMC/PBMC (2 x 10^6^ cells/well) were then stained with MR1-5-OP-RU tetramer at a dilution of 1:250 for 45 minutes at room temperature.^21^ Cells were then subsequently stained with an antibody cocktail of surface stains (listed in Supplemental Table 8) for 30 min at 4°C to allow non-CD3+ T cells or dead cells to be excluded. Cells were incubated in Human TruStain FcX (Biolegend^TM^) for 10 minutes at 4°C and then stained with TotalSeq-C hashtag antibodies and CITE-seq antibodies (Biolegend^TM^) listed in Supplemental Table 9 for 30 minutes at 4°C. Samples were then then sorted using a BD FACSAria^TM^ III Cell Sorter for MR1-5-OP-RU tetramer positive cells (representative gating strategy shown in Supplemental Figure 7). Sorted MR1T cells were then loaded in 10X Genomics^TM^ Chromium Next GEM Single Cell 5’ v2. Full protocol details of single-cell GEX, TCR and cell surface protein library preparation can be found from 10X genomics website (https://cdn.10xgenomics.com/image/upload/v1722286086/support-documents/CG000330_Chromium_Next_GEM_Single_Cell_5_v2_Cell_Surface_Protein_UserGuide_RevG.pdf). Finished libraries were then sent to Novogene Corporation Inc.^TM^ in Sacramento California for NovaSeq 6000 sequencing. MR1T cell clones were also sequenced using 10X Genomics^TM^ Chromium Next GEM Single Cell 5’ v2. Clones were either unstimulated at time of sequencing or were stimulated with PMA/Ionomycin for 3 hours prior to sequencing.

### Single-cell RNA-seq pre-processing

Raw sequence reads were processed using 10X Genomics Cell Ranger software (version 6.1.1). The resulting sequence data were aligned to the GRCh38 human genome. Cell demultiplexing used a combination of algorithms, including GMM-demux, demuxEM and BFF, implemented using the cellhashR package.^51–53^ Droplets identified as doublets (i.e. the collision of distinct sample barcodes) were removed from downstream analyses. We additionally performed doublet detection using DoubletFinder, and removed doublets from downstream analysis.^54^ Next, droplets were filtered based on UMI count (allowing 0-20,000/cell), and unique features (allowing 200-5000/cell). Additionally, we computed a per-cell saturation statistic for both RNA and antibody derived tags (ADT) data, defined as: 1 – (#UMIs / #Counts). This statistic provides a per-cell measurement of the completeness with which unique molecules are sampled per cell and has the benefit of being adaptable across diverse cell types. Data were filtered to require RNA saturation > 0.35. Analyses utilized the Seurat R package, version 4.2.^55^ Using standardized methods implemented in the Seurat R package, counts and UMIs were normalized across cells, scaled per 10,000 bases, and converted to log scale using the ‘NormalizeData’ function. These values were then converted to z-scores using the ‘ScaleData’ command. Highly variable genes were selected using the ‘FindVariableGenes’ function with a dispersion cutoff of 0.5. Principal components were calculated for these selected genes and projected onto all other genes using the ‘RunPCA’ and ‘ProjectPCA’ commands. Clusters of similar cells were identified using the Louvain method for community detection, and UMAP projections were calculated. CITE-seq data were CLR-normalized by lane, meaning raw count data are subset per lane, CLR normalization performed (as implemented in the Seurat R package, using margin = 1), using all QC-passing cells/lane. Normalization per-lane was performed to reduce batch effects.

### TCR sequence analysis

Raw sequence reads for gene expression and TCR enrichment were first processed using cellranger software, version 6.1.1 (10X Genomics). The raw clonotype calls produced by cellranger vdj were extracted from the comma-delimited outputs. Cells were demultiplexed and TCR calls were assigned to samples using custom software, made publicly available through the cellhashR package, with the GMM-Demux, demuxEM, and BFF algorithms.^51–53^ The clonotype data generated by cellranger were filtered to drop any cells where the TCR calls were lacking a CDR3 sequence, the clonotype was not marked as full-length, or the clonotype lacked a called V, J, or constant gene. Rows with chimeric V/J/C combinations (i.e. TRBV / TRAC) were filtered, with the exception that segments consisting of a TRDV/TRAC or TRAV/TRDC were permitted. These chimeric segments were classified according to the constant region chain. Data was analyzed in R studio using Seurat package.^56^

### Limited Dilution Cloning (LDC)

The protocol for LDC has been previously published.^57^ CBMC were stained with Propidium Iodide (Milteny Biotec), MR1-5-OP-RU tetramer (NIH tetramer core), and antibody cocktail of surface stains to sort out non-CD3+ cells (listed in Supplemental Table 8). Live, MR1/5-OP-RU tetramer positive cells were sorted and then cryopreserved. After thawing, the cells were plated in a 96-well plate in a limited dilution assay with 1.5x10^5^ irradiated PBMC (3000 cGray) and 3x10^4^ irradiated LCL (6000 cGray) along with αCD3 (30ng/ml) and IL-2 (2ng/ml) in RPMI 1640 supplemented with 10% heat inactivated human serum in 96 well round bottom plates. On day 5, the αCD3 was washed out by removal of half of the volume from the wells and replaced with additional IL-2 supplemented media. Media was changed for all wells every 2-3 days and replaced with IL-2 supplemented media. T cell “buttons” were evaluated for growth on Day 20 and selected buttons stained with the MR1-5-OP-RU tetramer, CD3 PeCy7 (clone SK7; Biolegend), CD4 FITC (clone SK3; Biolegend), CD8 APC Cy7 (clone SK1; Biolegend), and TRAV1-2 BV605 (clone IP26; Biolegend). T cell clones that remained MR1-5-OP-RU tetramer positive were re-expanded in T12.5 flasks with irradiated PBMC and LCL as well as RPMI 1640 media with 10% Human serum supplemented with αCD3 and IL-2 as above.^3^

### Clone phenotypic confirmation

Following LDC, isolated clones were analyzed phenotypically by flow cytometry to confirm retention of positive staining for the MR1-5-OP-RU tetramer and clonality. Clones were thawed in the presence of DNase, resuspended in 10% heat inactivated human serum with RPMI (Lonza) and counted. Viable cells (10^5^ cells/well) from each clone were plated in duplicate into a 96 well plate. Clones were washed in 2% fetal bovine serum (FBS) in phosphate buffered saline (PBS), following by staining with either MR1-5-OP-RU tetramer or MR1/6-FP tetramer at a dilution of 1:500 for 45 minutes. Following this, cells were stained with surface stains (Supplemental Table 10) for 30 minutes. Cells were washed in PBS and then resuspended in 1% paraformaldehyde and analyzed on a FACSymphony^TM^. Data was subsequently analyzed using FlowJo^TM^.

### Microbes and antigens

*E. coli, S. aureus, S. pneumoniae* were purchased from American Type Culture Collection (ATCC^TM^) and instructions for resuspension and growth were followed. Following growth of each bacteria, the concentration was calculated through serial dilutions and plating. Appropriate multiplicity of infection was determined through titrations using an adult-derived MAIT cell clone (D426 G11) to confirm adequate response to each bacterium. *M. tuberculosis* (Mtb) auxotroph mc^2^6206 (H37Rv ΔpanCD ΔleuCD)^36^ was as a kind gift from Bill Jacobs (MTA-IN19-168) and is modified such that it is only able to grow in the presence of externally supplemented pantothenate and leucine and thus can be handled in a biosafety level 2 setting. *M. smegmatis* is used frequently in our lab and two *M. smegmatis* MR1 ligands, photolumazine I (PL1) and deazalumazine (DZ), were previously generated and details of this have been previously published.^5^

### ELISpot analysis

Microbe and antigen responses from MR1T cell clones were determined using an enzyme-linked immunosorbent spot (ELISpot) assay for IFN-γ as described previously.^58^ Human DC were isolated from PBMC by plate adherence and cultured in RPMI+10% heat-inactivated (hi)-HuS with 30ng/ml GM-CSF (Immunex) and 10 ng/ml IL-4 (R&D Systems) as previously described.^59^ ELISpot plates were coated with anti-IFN-γ antibody (Mabtech, clone 1-D1K, Cat No. 3420-3-1000, used at 10 μg/ml). After overnight incubation at 4°C, ELISpot plates were washed three times with phosphate-buffered saline and then blocked with RPMI 1640 + 10% human serum for at least 1 hr. For experiments with Mtb, 5x10^5^ DC were plated in each well of an ultra-low adherence 24 well tissue culture plate. Mtb was grown fresh in 7H9 Middlebrook broth, supplemented with 50 μg/mL of leucine and 24 μg/mL of pantothenate, grown to an optical density of 0.5. 40ul (MOI 5.6), added and incubated overnight. DC (1x10^4^ cells^)^ were plated in the ELISpot wells in RPMI+10% HuS. For experiments with *E. coli, S. aureus, S. pneumoniae* and *M. smegmatis*, PL1, and DZ, DC (1x10^4^ cells) were plated in the ELISPOT wells in RPMI+10% HuS. The bacteria or antigens were added to the wells at concentrations or MOI indicated for each experiment and incubated for 1 hour. For all experiments, T cell clones (5000 cells) were added, and the plate was incubated overnight at 37°C. IFN-γ ELISpots were enumerated following development as previously described.^58^ Antibody blocking was performed using the anti-MR1 26.5 clone (OHSU antibody core) and an IgG2a isotype control (Biolegend, clone MOPC-173, Cat No. 400224) added at 5 μg/ml for 1 h prior to the addition of ligand. All samples were done as technical replicates in duplicate, and three independent biological replicates were done on separate days. Negative and positive controls were also performed using T cells and DC without antigen or with Phytohaemagglutin P (PHA, 10 μg/ml; Sigma Aldrich). Background spots seen in negative control were subtracted from the remainder of the plate. All ELISpot analysis and graphing were done in GraphPad Prism.

### Expression, refold and purification of MR1-β2m and MAIT TCR proteins

Human A-F7 (TRAV1-2/TRBV6-1),^38^ #6 (TRAV1-2-TRBV6-4),^38^ and CB964 A2 TCR (TRAV1-2-TRBV6-2) MAIT TCR proteins were refolded from inclusion bodies at 4°C for an overnight period using the refolding buffer that contained 0.1 M Tris pH 8.5, 6 M urea, 2 mM EDTA, 0.4 M L-arginine, 0.5 mM oxidized glutathione, and 5 mM reduced glutathione as previously reported.^38^ The wild-type or C-terminal cysteine mutated MR1-β2m was refolded at a 10x molar ratio of the investigated compound in the same refold buffer, as previously described.^39^ The refolded MR1-ligand and TCR proteins were purified using three sequential purification procedures: crude DEAE anion exchange, S200 15/60 size exclusion chromatography, and HiTrap-Q HP or MonoQ 10/100 GL anion exchange. The protein purity was assessed using sodium dodecyl sulfate–polyacrylamide gel electrophoresis (SDS-PAGE) and quantified using a NanoDrop™□ spectrophotometer.

### Surface Plasmon Resonance measurements

All Surface Plasmon Resonance (SPR) experiments were conducted at 25°C, in duplicate (n=3), on a BIAcore 3000 instrument using HBS buffer (10 mM HEPES-HCl pH 7.4, 150 mM NaCl, and 0.005% surfactant P20) as described previously.^60^ Biotinylated MR1-β2m-Ag was immobilized on SA-Chip (GE) with a surface density of ∼2,000 response units (RU). Various concentrations (0-200 µM) of two adult MAIT TCRs; A-F7 (TRAV1-2-TRBV6-1) and #6 (TRAV1-2-TRBV6-4), as well as the CB964 A2 TCR (TRAV1-2-TRBV6-2) were injected over the bound MR1-β2m-Ag at 5 µL/min and equilibrium data were collected. The final response was calculated by subtracting the response of the blank flow cell alone from the TCR-MR1-β2m-Ag complex. The SPR sensograms, equilibrium curves and steady state K_D_ values (µM) were prepared using GraphPad Prism 10.

### Crystallization, data Collection, structure determination, and analysis

Purified CB964 A2 TCR was mixed with MR1-β2m-5-OP-RU in a 1:1 molar ratio and then held on ice for two hours. The concentration of the mixture was 4-6 mg/ml. CB964 A2 TCR-MR1-5-OP-RU crystals were created at 20°C using the hanging-drop vapor diffusion method, as previously reported.^39^ The precipitant reservoir solution consisted of 100 mM Bis–Tris Propane (BTP; pH 6.0–6.7), 10–20% PEG3350, and 200 mM sodium acetate. The ternary complex crystals were grown for 6 weeks and harvested before being quickly soaked in reservoir solution containing 15% glycerol for cryoprotection and then flash-frozen in liquid nitrogen. At the Australian Synchrotron, X-ray diffraction data sets were collected at 100 K using the MX2 beamline. Diffraction data were processed using XDS^61^ and programs from the CCP4 suite^62^ and Phenix package.^63^ The ternary structure of CB964 A2 TCR-MR1-5-OP-RU was determined by molecular replacement using PHASER,^64^ where modified TCR-MR1 ternary complex (PDB: 6PUC) was used as a search model. After building the models in COOT^65^ and refining them iteratively with Phenix.refine,^63^ the models were validated with MolProbity.^66^ Supplemental Table 7 provides a summary of CB964 A2 TCR-MR1-5-OP-RU final crystallographic data. BSA calculations were performed using PISA^67^ and the contacts generated by the Contact program from the CCP4 suite.^62^ Molecular graphics visualisation was generated using PyMOL v2.5 (Schrödinger).

### Statistics

Statistical analysis was done in R using built-in statistical packages for all differential gene expression of single-cell gene expression data. Given the possibility of increased false discoveries with single-cell data due to mean expression bias,^68^ we transformed our single-cell gene expression data into pseudobulked gene expression prior to performing differential gene expression using the edgeR package.^69^ The only exception to this, was the analysis of CB participants’ TRAV1-2(+) MAIT cells compared to TRAV1-2(-) MR1T cells given the lower cell counts for this analysis. For this analysis we used a Wilcoxon test for statistical significance. For all analysis we considered a 2-log threshold change of more than 0.5 and a false discovery rate (FDR) or Bonferroni-corrected p-value of < 0.05 to be significant. For TCR analysis, data was first analyzed in R studio and then diversity and pairing preference data was extracted and analyzed in Prism. Statistics were performed in Prism using a t-test for significance, with p < 0.05 being considered significant.

## Data Availability

All R studio code used in the generation of this analysis is freely available and uploaded to github (https://github.com/kaindylan/Cordblood-Paper). All raw data was deposited to dbgap (accessions number: phs003810.v1.p1). The authors declare that the data supporting the findings of this study are available within the paper and its supplementary information files. The source data underlying the graphs in the paper can be found in the Supplementary Data. The accession number for the atomic coordinates of CB964 A2 TCR-MR1-5-OP-RU, along with associated structure factors, have been deposited at the protein databank (www.rcsb.org) with accession code 9MS0.

## Author Contribution

DK, GS, WA, JR, DML, and DAL contributed to the conception and/or design of the work. MC, GS, MN, DAL and DML contributed to clinical activities. DK, WA, JR, DML, and DAL raised grants to fund the research. DK, MC, GS, GM, WA, KCYP, CM, GB, KR, MN, DML, JR, BB, and DAL substantially contributed to the acquisition, analysis, or interpretation of data and drafting of the manuscript. All authors substantially contributed to revising and critically reviewing the manuscript for important intellectual content. All authors approved the final version of this manuscript to be published and agree to be accountable for all aspects of the work.

## Acknowledgements

We would like to thank the participants who gave time and dedication to this health research as well as Erin Merrifield, Department of Pediatrics, OHSU, and Dr. Marielle Gold for their contributions to this study. We thank the staff at the Monash Macromolecular Crystallization Facility. This research was undertaken in part using the MX2 beamline at the Australian Synchrotron, part of ANSTO, and used the Australian Cancer Research Foundation (ACRF) detector.

## Supplemental

**Supplemental Figure 1:**
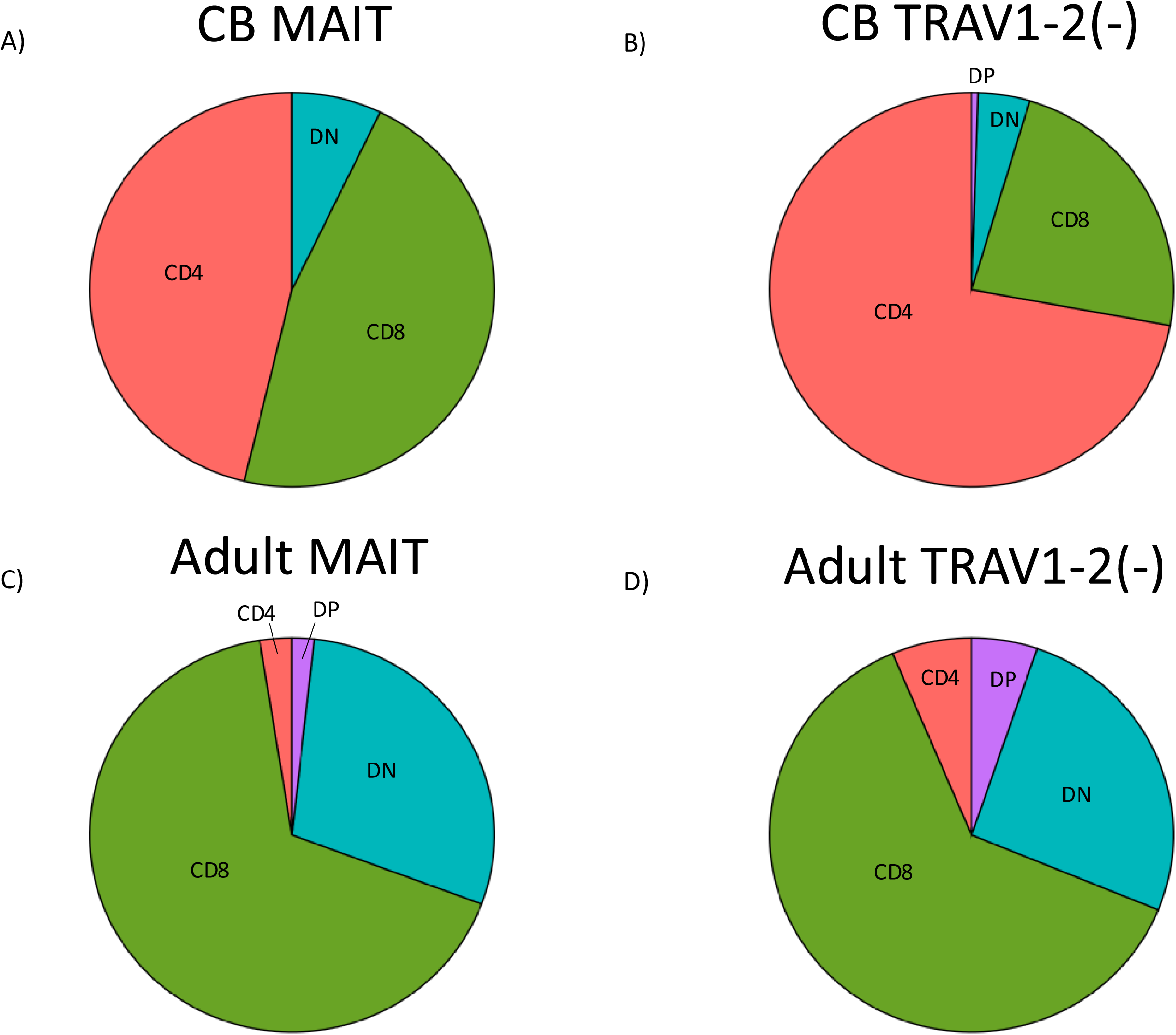
CD4 and CD8 Protein Expression. a-b) CB MR1T cell expression of CD8 and CD4 (based on CITE-seq cell surface expression), separated by TRAV expression (based on TCR sequencing). c-d) Adult MR1T cell expression of CD8 and CD4 (based on CITE-seq cell surface expression), separated by TRAV expression (based on TCR sequencing). DP = double positive, DN = double negative.

**Supplemental Figure 2:**
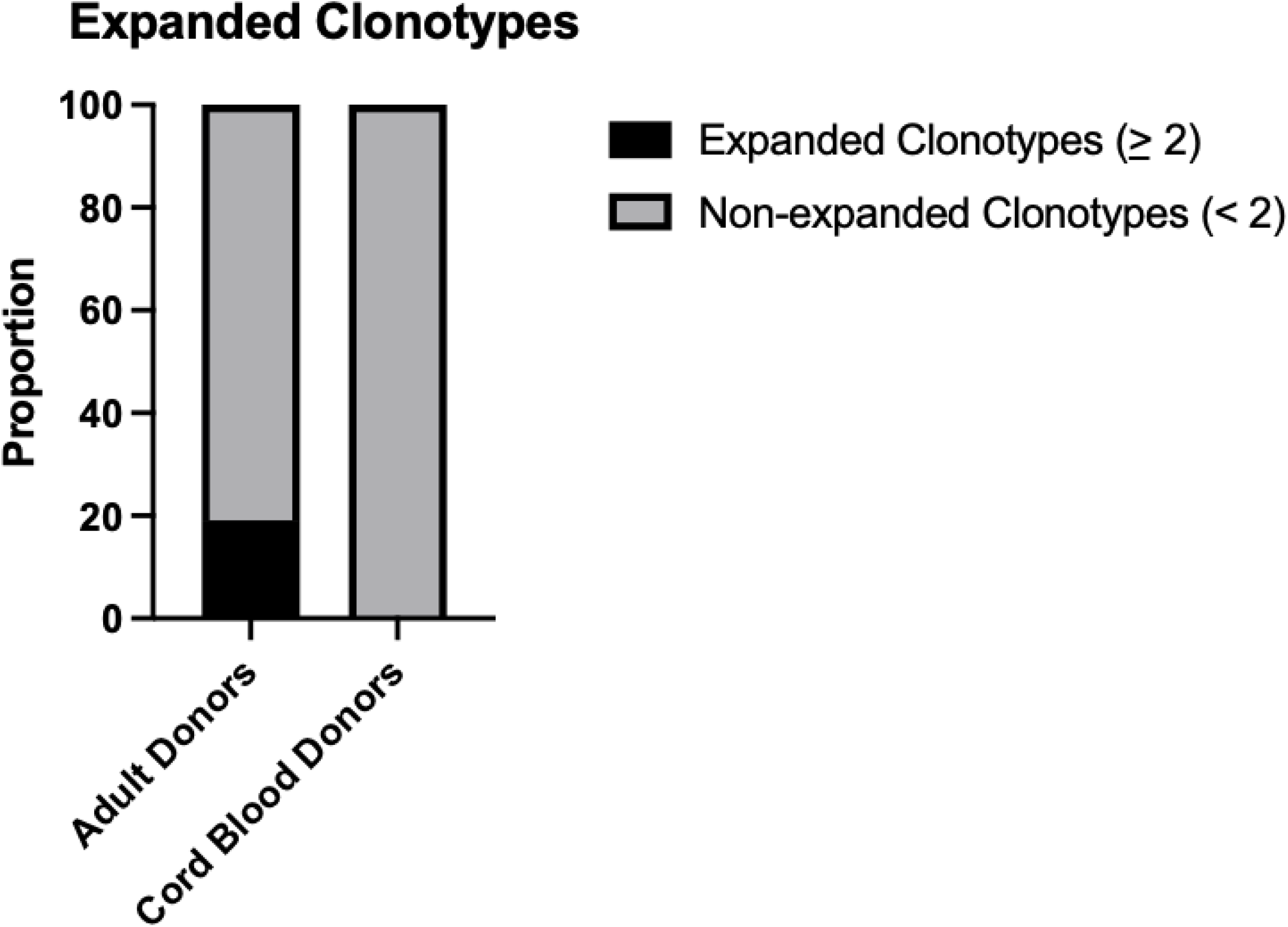
Proportion of expanded clones: Proportion of cells from adult or CB participants with at least one other cell with an identical CDR3.

**Supplemental Figure 3:**
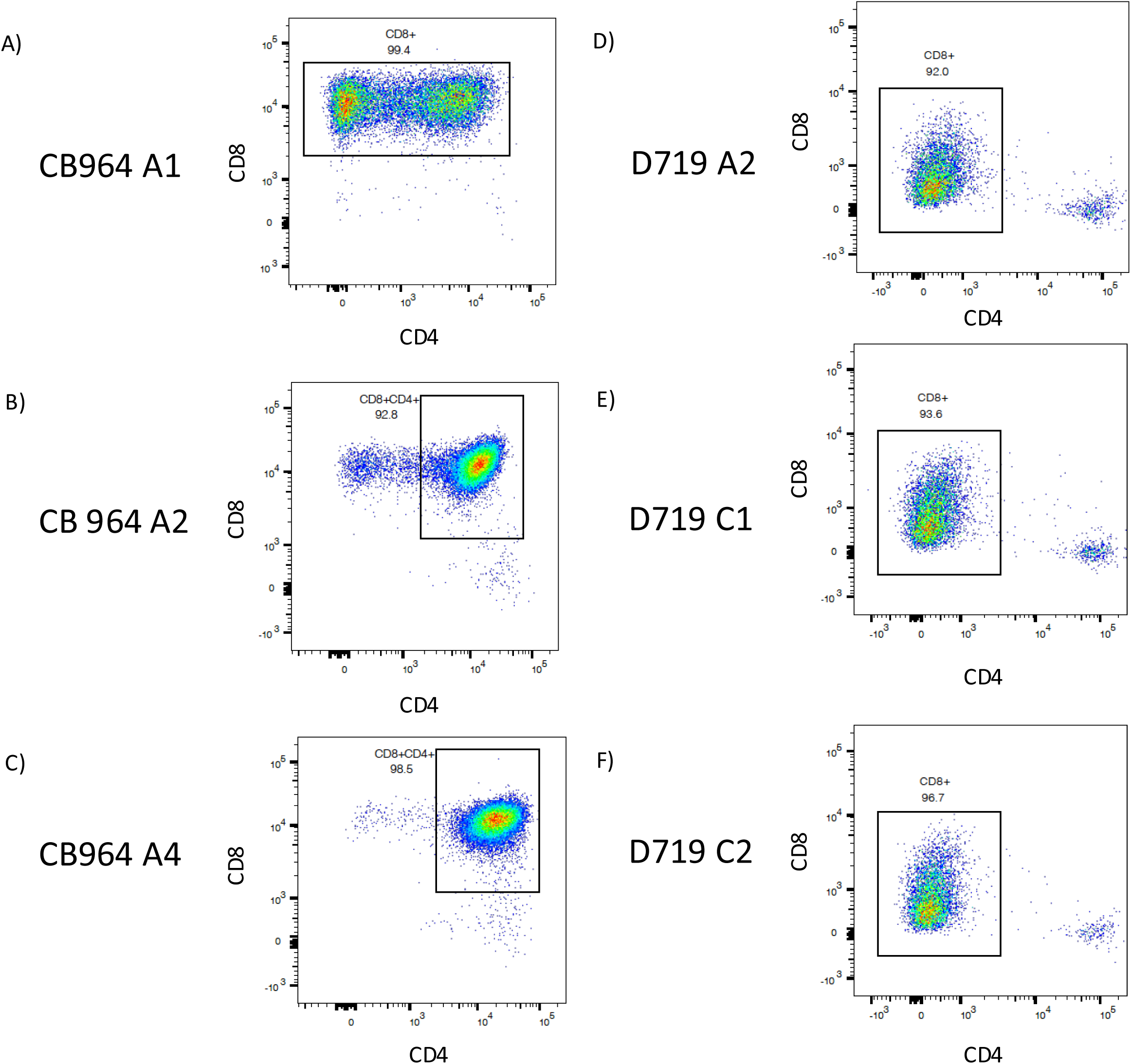
CD8 and CD4 expression of CB and adult MR1T cell clones. a-c) CD4, CD8, TRAV1-2 and tetramer staining of CB MR1T cell clones by flow cytometry d-f) CD4, CD8, TRAV1-2 and tetramer staining of adult MR1T cell clones by flow cytometry

**Supplemental Figure 4:**
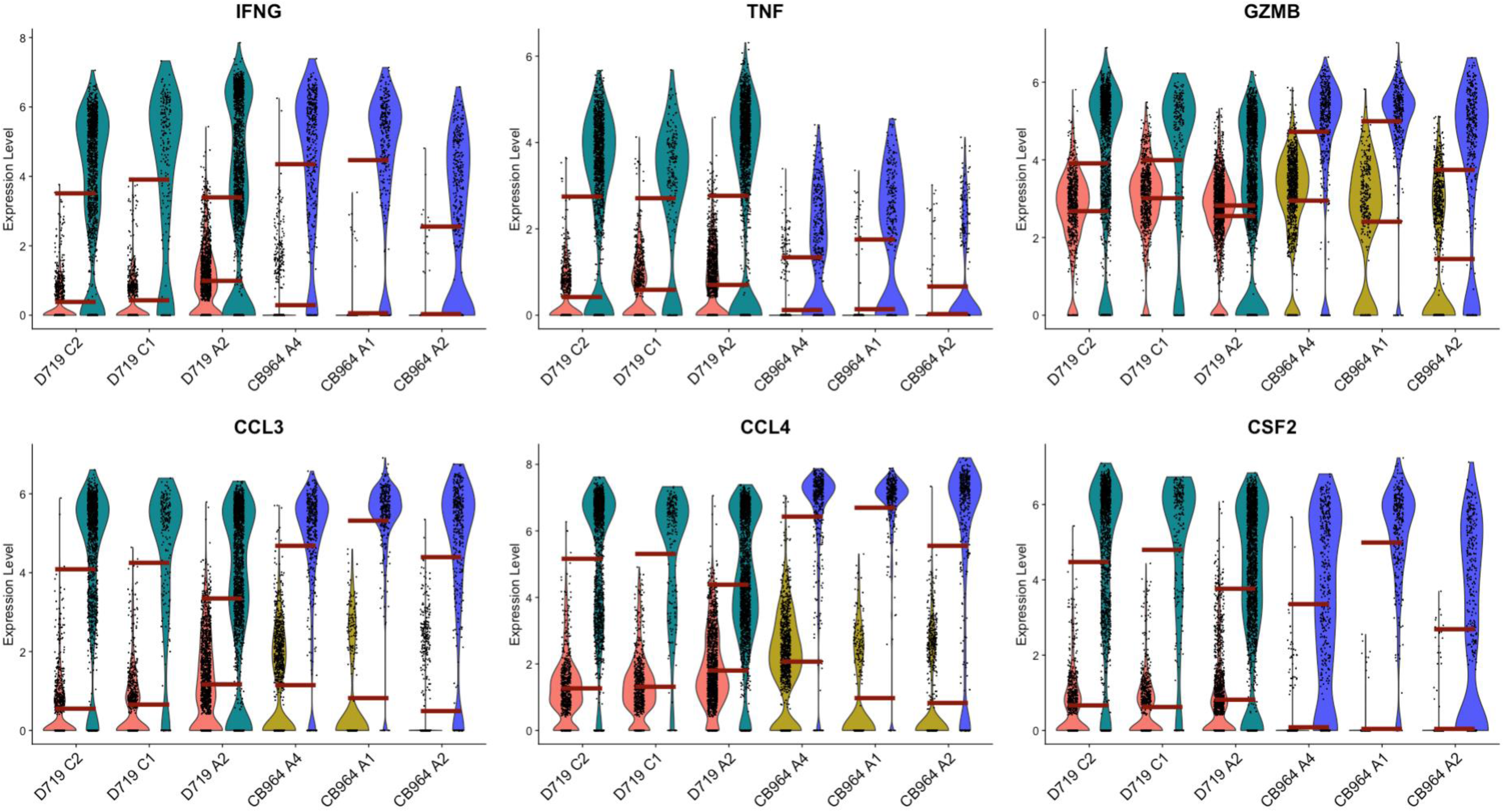
CB MR1T cell clones can produce typical inflammatory/cytotoxic cytokines by gene expression at individual clone level. Violin plot of gene expression of common inflammatory/cytotoxic genes from stimulated (PMA/Ionomycin) and unstimulated MR1T cell clones from adult compared to CB donors broken out by each individual clone.

**Supplemental Figure 5:**
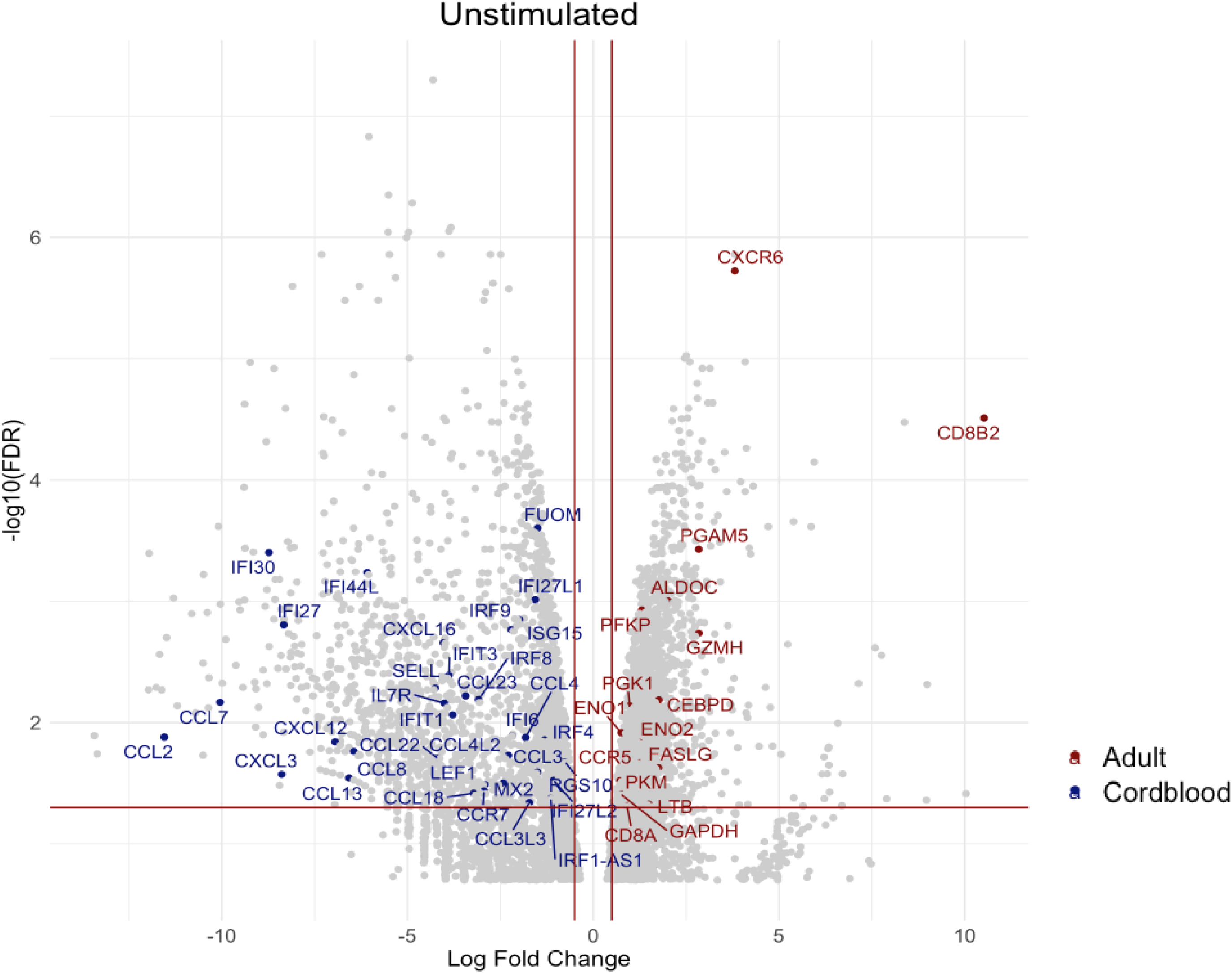
Differential gene expression of CB and adult MR1T cell clones. Volcano plot of differentially expressed genes of adult compared to CB MR1T cell clones at unstimulated state. Horizontal red line represents Bonferroni-adjusted p-value of 0.05, and vertical red lines represent log fold change of 0.5 or -0.5.

**Supplemental Figure 6:**
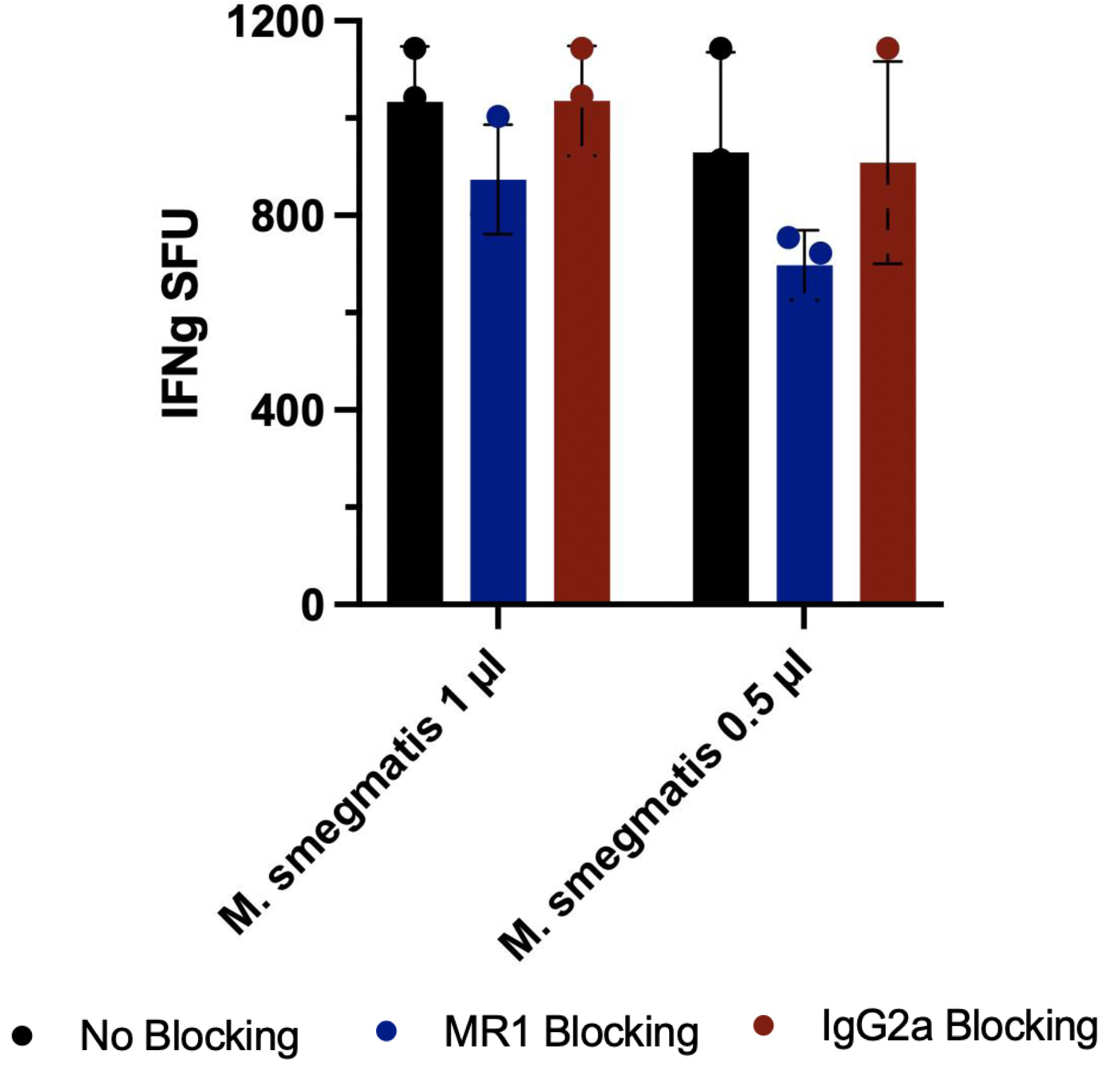
Titration of *Mycobacterium smegmatis*. IFNγ ELISPOT of adult MR1T cell clone (G11) with lower concentrations of *M. smegmatis.* Responses without blocking, anti-MR1 blocking antibody or isotype control blocking antibody are represented.

**Supplemental Figure 7:**
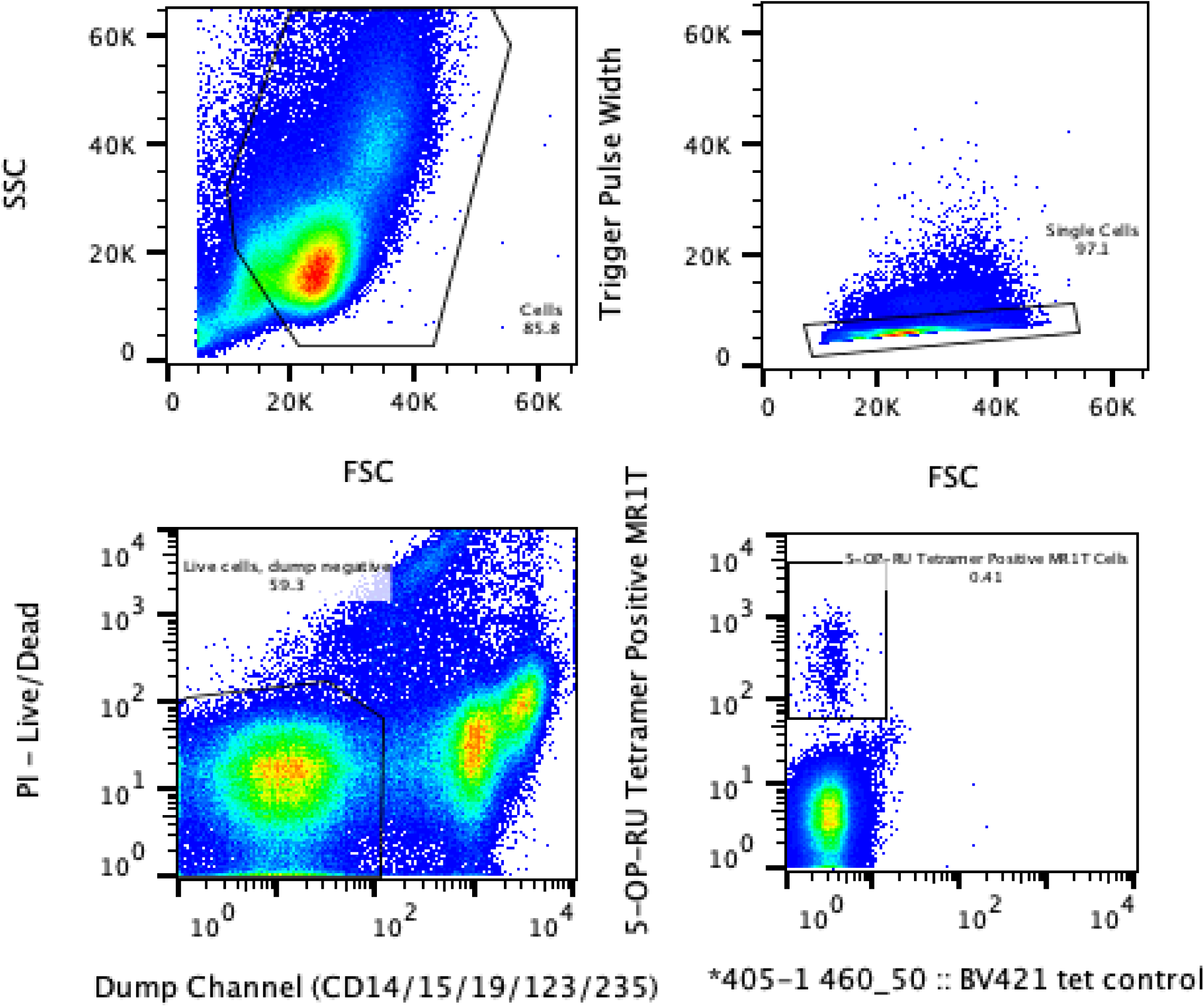
Representative FACS Sorting Strategy. Live MR1/5-OP-RU positive cells were sorted based on PI negativity, negative staining (dump channel) for CD14, CD15, CD19, CD123, CD235ab (PeCy7), and MR1/5-OP-RU tetramer positive.

## Cord Blood Paper References

1 World Health Organization. Global Report on the Epidemiology and Burden of Sepsis. 2020.

2 World Health Organization. Global Tuberculosis Report. 2021

3 Gold MC, Cerri SC, Smyk-Pearson S, et al. Human mucosal associated invariant T cells detect bacterially infected cells. PLoS Biol. 2010;8(6): e1000407

4 Le Bourhis L, Martin E, Peguillet I, et al. Antimicrobial activity of mucosal-associated invariant T cells. Nat Immunol. 2010;11:701–708

5 Harriff M, McMurtry C, Floyd CA, et al. MR1 displays the microbial metabolite driving selective MR1-resitricted T cell receptor usage. Science Immunology. 2018;3(25):eaao2556

6 Meermeier EW, Laugel BF, Sewell AK, et al. Human TRAV1-2-negative MR1-restricted T cells detect *S. pyogenes* and alternatives to MAIT riboflavin-based antigens. Nature communications. 2016;7:12506

7 McInerney MP, Awad W, Souter MNT and Purcell AW. MR1 presents vitamin B6-related compounds for recognition by MR1-reactive T cells. PNAS. 2024;121(49):e2414792121

8 Ito E, Inuki S, Takahashi M, et al. Sulfated bile acid is a host-derived ligand for MAIT cells. Science Immunology. 2024;9(91):ade6924

9 Vacchini A, Chancellor A, Yang Q, et al. Nucleobase adducts bind MR1 and stimulate MR1-restricted T cells. Sci Immunol. 2024;9(25):eadn0126

10 Keller AN, Eckle SBG, Xu W, et al. Drugs and drug-like molecules can modulate the function of mucosal-associated invariant T cells. Nat Immunol. 2017;18(4):402–411

11 Lepore M, Kalinicenko A, Colone A, et al. Parallel T-cell cloning and deep sequencing of human MAIT cells reveal stable oligoclonal TCR beta repertoire. Nat Commun. 2014;5:3866

12 Walker LJ, Kang YH, Smith MO, et al. Human MAIT and CD8alphaalpha cells develop from a pool of type-17 precommitted CD8+ T cells. Blood. 2012;119(2):422–33

13 Fergusson JR, Smith KE, Fleming VM, et al. CD161 defines a transcriptional and functional phenotype across distinct human T cell lineages. Cell Rep. 2014;9(3):1075–88

14 Sharma PK, Wong EB, Napier RJ, et al. High expression of CD26 accurately identifies human bacteria-reactive MR1-restricted MAIT cells. Immunology. 2015;145(3):443–53

15 Gherardin NA, Keller AN, Woolley RE, et al. Diversity of T cells Restricted by the MHC Class I-Related Molecule MR1 Facilitates Differential Antigen Recognition. Immunity. 2016;44(1):32–45

16 Kjer-Nielsen L, Patel O, Corbett AJ, et al. MR1 presents microbial vitamin B metabolites to MAIT cells. Nature. 2012;491(7426):717–23

17 Georgel P, Radosavljevic M, Macquin C, and Bahram S. Ten non-conventional MHC class I MR1 molecule controls infection by *Klebsiella pneumoniae* in mice. Mol Immunol. 2011;48:769–775

18 Riffelmacher T, Murray MP, Wientjens C, et al. Divergent metabolic programs control two populations of MAIT cells that protect the lung. Nature Cell Biology. 2023;25:877–891

19 Lepore M, Kalinichenko A, Calogero S, et al. Functionally diverse human T cells recognize non-microbial antigens presented by MR1. eLife. 2017;6:e24476

20 Crowther MD, Dolton G, Legit M, et al. Genome-wide CRISPR-Cas9 screening reveals ubiquitous T cell cancer targeting via the monomorphic MHC class I-related protein MR1. Nat Immunol. 2020;21(2):178–185

21 Swarbrick GM, Gela A, Cansler ME, et al. Postnatal Expansion, Maturation, and Functionality of MR1T Cells in Humans. Frontiers in Immunology. 2020;11:5566995

22 Constantinides MG, Link VM, Tamoutounour S, et al. MAIT cells are imprinted by the microbiota in early life and promote tissue repair. Science. 2019;366(6464):eaax6624

23 Carmona Lab. UCell. Accessed May 15^th^, 2024 <https://github.com/carmonalab/UCell>

24 Suliman S, Kjer-Nuelsen L, Iwany SK, et al. Dual TCR-alpha expression on MAIT cells as a potential confounder of TCR interpretation. J Immunol. 2022;208(6):1389–1395

25 Szabo PA, Levitin HM, Miron M, et al. Single-cell transcriptomics of human T cells reveals tissue and activation signatures in health and disease. Nat Comms. 2019;10:4706

26 Ingram RJ, Isaacs JD, Kaur G, et al. A role of cellular prion protein in programming T-cell cytokine responses in disease. FASEB J. 2009;23(6):1672–84

27 Gingras S, Pelletier S, Boyd K, and Ihle JN. Characterization of a Family of Novel Cysteine-Serine-Rich Nuclear Proteins (CSRNP). PLoS One. 2007;2(8):e808

28 SH2D2A. Bethesda (MD): National Library of Medicine (US), National Center for Biotechnology Information; 2004 – [cited 2024, Jan 14^th^]. Available from: https://www.ncbi.nlm.nih.gov/gene/

29 Meixner A, Karreth F, Kenner L, and Wagner EF. JunD regulates lymphocyte proliferation and T helper cell cytokine expression. The EMBO Journal. 2004;23:1325–1335

30 GNA15. Bethesda (MD): National Library of Medicine (US), National Center for Biotechnology Information; 2004 – [cited 2024, Jan 14^th^]. Available from: https://www.ncbi.nlm.nih.gov/gene/

31 Kouwaki T, Okamoto M, Tsukamoto H, et al. Zyxin stabilizes RIG-I and MAVS interactions and promotes type I interferon response. Sci Rep. 2017;7(1):11905

32 Fonseca R, Burn TN, Gandolfo LC, et al. Runx3 drives a CD8^+^ T cell tissue residency program that is absent in CD4^+^ T cells. Nature Immunology. 2022;23:1236–1245

33 Savinko T, Guenther C, Uotila LM, et al. Filamin A Is Required for Optimal T Cell Integrin-Mediated Force Transmission, Flow Adhesion, and T Cell Trafficking. The Journal of Immunology. 2018;200(9):3109–3116

34 Hengel RL, Thaker V, Pavlick MV, et al. Cutting Edge: L-Selectin (CD62L) Expression Distinguishes Small Resting Memory CD4^+^ T Cells That Preferentially Respond to Recall Antigen. The Journal of Immunology. 2003;170(1):28–32

35 Cassidy FC, Kedia-Mehta N, Bergin R and Hogan AE. Glycogen-fueled metabolism supports rapid mucosal-associated invariant T cell responses. PNAS. 2023;120:e2300566120

36 Mouton JM, Heunis T, Dippenaar A, et al. Comprehensive Characterization of the Attenuated Double Auxotroph Mycobacterium tuberculosis ΔleuDΔpanCD as an Alternative to H37Rv. Front Microbiol. 2019;10:1922

37 Awad W, Ler GJM, Xu W, et al. The molecular basis underpinning the potency and specificity of MAIT cell antigens. Nat Immunol. 2020;21:400–411

38 Eckle SBG, Birkinshaw RW, Kostenko L, et al. A molecular basis underpinning the T cell receptor heterogeneity of mucosal-associated invariant T cells. JEM. 2014;211(8):1585–1600

39 Corbett AJ, Eckle SBG, Birkinshaw RW, et al. T-cell activation by transitory neo-antigens derived from distinct microbial pathways. Nature. 2014;509(7500):361–5

40 Awad W, Ciacchi L, McCluskey J et al. Molecular insights into metabolite antigen recognition by mucosal-associated invariant T cells. Current Opinion in Immunology. 2023;83:102351

41 Leeansyah E, Loh L, Nixon DF, and Sandberg JK. Acquisition of innate-like microbial reactivity in mucosal tissues during fetal MAIT-cell development. Nat Comms. 2014;5:3143

42 Garner LC, Amini A, FitzPatrick MEB, et al. Single-cell analysis of human MAIT cell transcriptional, functional and clonal diversity. Nat Immunology. 2023(24):1565–1578

43 Koay HF, Gherardin NA, Enders A, et al. A three-stage intrathymic development pathway for the mucosal-associated invariant T cell lineage. Nature Immunology. 2016;17(11):1300–1311

44 Koay HF, Su S, Amann-Zalcenstein D, et al. A divergent transcriptional landscape underpins the development and functional branching of MAIT cells. Science Immunology. 2019;4(41):eaay6039

45 Davis MM, and Boyd SD. Recent progress in the analysis of αβ T cell and B cell receptor repertoires. Current Opinion in Immunology. 2019;59:109–114

46 Gold MC, McLaren JE, Reistetter JA, et al. MR1-restricted MAIT cells display ligand discrimination and pathogen selectivity through distinct T cell receptor usage. JEM. 2014;211(8):1601–1610

47 Narayanan GA, McLaren JE, Meermeier EW, et al. The MAIT TCRβ chain contributed to discrimination of microbial ligand. Immunol Cell Biol. 2020;98(9):770–781

48 Dias J, Leeansyah E, and Sandbery JK. Multiple layers of heterogeneity and subset diversity in human MAIT cell response to distinct microorganisms and to innate cytokines. Proc Natl Acad Sci USA. 2017;114(27):E5434–E5443

49 Chengalroyen MD, Oketada N, Worley A, et al. Disruption of riboflavin biosynthesis in mycobacteria establishes 5-amino-6-D-ribitylaminouracil (5-A-RU) as key precursor of MAIT cell agonists. bioRxiv. 2024.10.03.616430

50 Pieren DKR, Boer MC, and de Wit J. The adaptive immune system in early life: The shift makes it count. Front Immunol. 2022;13:1031924.

51 Xin H, Lian Q, Jiang Y, et al. GMM-Demux: sample demultiplexing, multiplet detection, experiment planning, and novel cell-type verification in single cell sequencing. Genome Biol. 2020;21(188):1–35

52 Boggy G, McElfresh GW, Mahyari E, et al. BFF and cellhashR: analysis tools for accurate demultiplexing of cell hashing data. Bioinformatics. 2022;38(10):2791–2801

53 Gaublomme JT, Li B, McCabe C, et al. Nuclei multiplexing with barcoded antibodies for single-nucleus genomics. Nature Communications. 2019;10(2907):1–8

54 McGinnis CS, Murrow LM, and Gartner ZJ. DoubletFinder: Doublet Detection in Single-Cell RNA Sequencing Data Using Artificial Nearest Neighbors. Cell Syst. 2019;8(4):329–337

55 Butler A, Hoffman P, Smibert P, et al. Integrating single-cell transcriptomic data across different conditions, technologies, and species. Nature Biotechnology. 2018;26:411–420

56 Seurat 4.3.0. Seurat – Guided Clustering Tutorial. Accessed June 30^th^, 2024 <https://satijalab.org/seurat/articles/pbmc3k_tutorial.html>

57 Cansler M, Null M, Meerimeier E, et al. Generation of MR1-Restricted T Cell Clones by Limiting Dilution Closing of MR1 Tetramer^+^ cells. Methods Mol Biol. 2020;2098:219–235

58 Heinzel AS, Grotzke JE, Lines RA, et al. HLA-E-dependent presentation of Mtb-derived antigen to human CD8+ T cells. J Exp Med. 2002;196:1473–1481

59 Harriff MJ, Cansler ME, Toren KG, et al. Human Lung Epithelial Cells Contain Mycobacterium tuberculosis in a Late Endosomal Vacuole and Are Efficiently Recognized by CD8^+^ T Cells. PLOS ONE. 2014;9(5):e97515

60 Patel O, Kjer-Nielsen L, Le Nours J, et al. Recognition of vitamin B metabolites by mucosal-associated invariant T cells. Nat Commun. 2013;2142(4):3142

61 Kabsch W. Integration, scaling, space-group assignment and post-refinement. Acta crystallographica Section D, Biological crystallography. 2010;66:133–144

62 Winn MD, Ballard CC, Cowtan KD, et al. Overview of the CCP4 suite and current developments. Acta crystallographica Section D, Biological crystallography. 2011;67:235–242

63 Adams PD, Afonine PV, Bunkoczi G, et al. PHENIX: a comprehensive Python-based system for macromolecular structure solution. Acta crystallographica Section D, Biological crystallography. 2010;66:213–21

64 McCoy AJ. Solving structures of protein complexes by molecular replacement with Phaser. Acta crystallographica Section D, Biological crystallography. 2007;63:32–41

65 Emsley P, and Cowtan K. Coot: model-building tools for molecular graphics. Acta crystallographica Section D, Biological crystallography. 2004;60:2126–2132

66 Chen VB, Arendall WB, Headd JJ, et al. MolProbity: all-atom structure validation for macromolecular crystallography. Acta crystallographica Section D, Biological crystallography. 2010;66:12–21

67 Krissinel E, and Henrick K. Inference of macromolecular assemblies from crystalline state. J Mol Biol. 2007;372:774–797

68 Squair JW, Gautier M, Kathe C, et al. Confronting false discoveries in single-cell differential expression. Nat Comms. 2021;12:5692

69 Robinson MD, McCarthy DJ, and Smyth GK. edgeR: a Bioconductor package for differential expression analysis of digital gene expression data. Bioinformatics. 2009;26(1):139–140

