## Supplemental Table 6 and 7 for "Human Neonatal MR1T Cells Have Diverse TCR Usage, are Less Cytotoxic and are Unable to Respond to Many Common Childhood Pathogens"

**Supplemental Table 6. Diffraction data collection and refinement statistics**

|  | Cord A2 TCR-MR1-5-OP-RU |
| --- | --- |
| Wavelength (Å) | 0.9537 |
| Resolution range (Å) | 46.84 - 2.52 (2.59 - 2.52) |
| Space group | P 21 21 21 |
| Unit cell  a, b, c (Å)  α, β, γ (°) | 62.526 114.533 141.3  90 90 90 |
| Total reflections | 69666 (5720) |
| Unique reflections | 34953 (2871) |
| Multiplicity | 2.0 (2.0) |
| Completeness (%) | 99.57 (99.65) |
| Mean I/sigma(I) | 10.20 (2.04) |
| Wilson B-factor | 41.61 |
| R-merge | 0.0467 (0.359) |
| R-meas | 0.06605 (0.5077) |
| R-pim | 0.0467 (0.359) |
| CC1/2 | 0.997 (0.768) |
| CC* | 0.999 (0.932) |
| R-work | 0.1827 (0.2765) |
| R-free | 0.2087 (0.3104) |
| Number of non-hydrogen atoms | 6708 |
| macromolecules | 6314 |
| ligands | 22 |
| solvent | 372 |
| Protein residues | 791 |
| RMSD (bonds) (Å) | 0.116 |
| RMSD (angles) (°) | 2.37 |
| Ramachandran favored (%) | 97.29 |
| Ramachandran allowed (%) | 2.58 |
| Ramachandran outliers (%) | 0.13 |
| Rotamer outliers (%) | 4.33 |
| Average B-factor | 47.41 |
| macromolecules | 47.46 |
| ligands | 28.16 |
| solvent | 47.65 |

Statistics for the highest-resolution shell are shown in parentheses.

**Supplemental Table 7. Contacts of A2 TCR with MR1-5-OP-RU**

| **TCR gene** | **TCR residues** | **MR1 residues** | **Bond type** |
| --- | --- | --- | --- |
| **CDR1⍺** | Gly^28^ | Glu^160^ | VDW |
|  | Phe^29^ | Glu^160^ | VDW |
|  | Phe^29-N^ | Glu^160-Oε2^ | H-bond |
|  | Phe^29-O^ | Asn^155-Nδ2^, Glu^160-Oε2^ | H-bond |
|  | Asn^30^ | Tyr^152^, Trp^156^, Glu^160^ | VDW |
| **CDR2⍺** | Tyr^48^ | His^148^, Tyr^152^ | VDW |
|  | Val^50^ | Leu^151^, Tyr^152^, Asn^155^ | VDW |
|  | Leu^51^ | Leu^151^, Lys^154^ | VDW |
|  | Glu^55^ | His^148^ | VDW |
| **FW⍺** | Arg^66^ | Asn^155^, Glu^159^ | VDW |
|  | Arg^66-Nη1^ | Asn^155-Oδ1^ | H-bond |
| **CDR3⍺** | Ser^93^ | Tyr^62^, Glu^160^, Trp^164^ | VDW |
|  | Ser^93-O^ | Tyr^62-OH^ | H-bond |
|  | Ser^94-O^ | Arg^61-Nε^, Arg^61-Nη2^ | H-bond |
|  | Ser^94^ | Arg^61^, Leu^65^ | VDW |
|  | Tyr^95^ | Arg^61^, Tyr^62^, Leu^65^, Tyr^152^, Trp^156^ | VDW |
|  | Tyr^95-O^ | Arg^61-Nη2^ | H-bond |
|  | Tyr^95-OH^ | Tyr^152-OH^, Trp^156-Nε1^ | H-bond |
| **CDR1β** | Tyr^31^ | Gly^68^ | VDW |
| **CDR2β** | Tyr^48-OH^ | Arg^61-Nη1^ | H-bond |
|  | Tyr^48^ | Arg^61^, Gln^64^ | VDW |
|  | Val^50^ | Gln^64^ | VDW |
| **FWβ** | Ala^56^ | Gln^64^ | VDW |
| **CDR3β** | Thr^97-Oɣ1^ | Trp^69-Nε1^ | H-bond |
|  | Thr^97^ | Leu^65^ | VDW |
|  | Asn^98-Oδ1^ | Glu^149-Oε2^ | H-bond |
|  | Asn^98-Nδ2^ | Glu^149-Oε2^ | H-bond |
|  | Asn^98^ | Glu^149^, Tyr^152^ | VDW |
|  | Tyr^99-OH^ | His^148-Nδ1^ | H-bond |
|  | Tyr^99^ | His^148^ | VDW |

- Atomic contacts determined using the *CONTACT* program of the CCP4i package with cutoff of 4 Å.
- Hydrogen bond interactions are defined as contact distances of 3.5 Å or less.
- Van der Waals (VDW) interactions are defined as non-hydrogen bond contact distances of less than 4 Å.
- R: 5-OP-RU
