## Supplemental Table 8 to 10 for "Human Neonatal MR1T Cells Have Diverse TCR Usage, are Less Cytotoxic and are Unable to Respond to Many Common Childhood Pathogens"

**Cord Blood Paper Supplemental Tables**

Supplemental Table 8: Reagents used for FACS sorting

| **Reagent** | **CAT** | **Company** |
| --- | --- | --- |
| CD14 PeCy7 | 301814 | Biolegend |
| CD15 PeCy7 | 301924 | Biolegend |
| CD19 PeCy7 | 302216 | Biolegend |
| CD123 PeCy7 | 306010 | Biolegend |
| CD235ab PeCy7 | 306620 | Biolegend |
| Proprium Iodide Solution | 130-093-233 | Miltenyi Biotec |

Supplemental Table 9: Reagents used for 10X Genomics® Staining

| **Reagent** | **CAT** | **Company** |
| --- | --- | --- |
| TotalSeq Hashtag 1 | 394661 | Biolegend |
| TotalSeq Hashtag 2 | 394663 | Biolegend |
| TotalSeq Hashtag 3 | 394665 | Biolegend |
| TotalSeq Hashtag 4 | 394667 | Biolegend |
| TotalSeq Hashtag 5 | 394669 | Biolegend |
| TotalSeq CD4 | 300567 | Biolegend |
| TotalSeq CD8 | 344753 | Biolegend |
| TotalSeq TCR Vα7.2 | 351735 | Biolegend |
| TotalSeq CD26 | 302722 | Biolegend |
| TotalSeq CD161 | 339947 | Biolegend |
| TotalSeq CD197 (CCR7) | 353251 | Biolegend |
| TotalSeq CD279 (PD-1) | 329963 | Biolegend |
| TotalSeq CD45RA | 304163 | Biolegend |
| Human TruStain FcX | 422302 | Biolegend |

Supplemental Table 10: Reagents used for MR1T cell clone flow cytometry

| **Reagent** | **CAT** | **Company** |
| --- | --- | --- |
| CD8 APC Cy7 | 344714 | Biolegend |
| CD3 BV421 | 317344 | Biolegend |
| CD4 BUV737 | 612748 | BD Horizon |
| CD26 FITC | 302704 | Biolegend |
| CD161 PeCy7 | 339918 | Biolegend |
| TRAV1-2 PerCPCy55 | 351710 | Biolegend |
| Live Dead Aqua | L34966 | Thermo Fischer Scientific |
